# Prenatal Methadone Exposure Disrupts Behavioral Development and Alters Motor Neuron Intrinsic Properties and Local Circuitry

**DOI:** 10.1101/2020.09.25.314013

**Authors:** Gregory G. Grecco, Briana Mork, Jui Yen Huang, Corinne E. Metzger, David L. Haggerty, Kaitlin C. Reeves, Yong Gao, Hunter Hoffman, Simon N. Katner, Andrea R. Masters, Cameron W. Morris, Erin A. Newell, Eric A. Engleman, Anthony J. Baucum, Jieun Kim, Bryan K. Yamamoto, Matthew R. Allen, Yu-Chien Wu, Hui-Chen Lu, Patrick L. Sheets, Brady K. Atwood

## Abstract

Despite the rising prevalence of methadone treatment in pregnant women with opioid use disorder, the effects of methadone on neurobehavioral development remain unclear. We developed a translational mouse model of prenatal methadone exposure (PME) that resembles the typical pattern of opioid use by pregnant women who first use oxycodone then switch to methadone maintenance pharmacotherapy, and subsequently become pregnant while maintained on methadone. We investigated the effects of PME on physical development, sensorimotor behavior, and motor neuron properties using a multidisciplinary approach of physical, biochemical, and behavioral assessments along with brain slice electrophysiology and in vivo magnetic resonance imaging. PME produced substantial impairments in offspring physical growth, activity in an open field, and sensorimotor milestone acquisition which were associated with alterations in motor neuron functioning and connectivity. The present study adds to the limited work examining PME by providing a comprehensive, translationally relevant characterization of how PME disrupts offspring development.

## INTRODUCTION

Pregnant women and their developing fetuses represent a vulnerable population severely impacted by the opioid crisis. From 1999-2014, the annual prevalence of opioid use disorder (OUD) in pregnant women at delivery increased 333%^1^. Mirroring this rise, neonatal opioid withdrawal syndrome (NOWS) increased from 1.2 to 5.8 per 1,000 hospital births from 2000 to 2012 with some geographical regions near 33 cases per 1,000^2,3^. Opioid maintenance therapies, such as methadone and buprenorphine, continue to represent the first-line treatments for pregnant women with OUD because these therapies benefit overall maternal-fetal outcomes at parturition^4^. However, it is unknown exactly how prenatal methadone exposure (PME) impacts the brain and behavioral development as these infants mature.

Prenatal opioid exposure is associated with smaller head circumferences^5^, lower brain volumes^6,7^, and deficits in white matter microstructure^8^, although these studies are limited by small sample sizes making it difficult to control for confounding variables. A recent, large study using data from the Adolescent Brain Cognitive Development study reported reduced motor cortical volumes and surface area in children with prenatal opioid exposure^7^. These differences remained significant when controlling for multiple additional factors (e.g. socioeconomic factors and prenatal alcohol/tobacco), indicating opioid exposure *in utero* may specifically disrupt motor cortex development and potentially impact motor behavior^7^. Prior studies revealed poorer motor performance in children exposed to opioids prenatally suggesting that prenatal opioid exposure produces long-lasting changes in both motor behavior and in brain regions associated with motor functioning^9,10^. However, clinical studies are often complicated by significant environmental variations which are difficult to control such as the extent of prenatal care or additional prenatal substance exposures which also impact neurodevelopmental outcomes of children^11^. Therefore, a translationally relevant animal model of prenatal opioid exposure is desperately needed to examine the neurological and behavioral consequences of opioid exposure in the absence of confounding variables.

Animal models of prenatal opioid exposure have demonstrated deficits in several sensorimotor milestones^12-14^, but the mechanisms underlying disrupted sensorimotor development are unclear. Preclinical studies suggest that prenatal opioid exposure may prevent normal neuronal development^15,16^ and disturb myelination^17^ which could underlie the aberrant neurobehavioral development. Unfortunately, the translational value of these rodent studies is limited by models that do not adequately model the majority of human prenatal opioid exposure cases. For instance, a large proportion of studies initiate opioid delivery around mid-gestation or later and often utilize morphine, even though methadone and buprenorphine account for a growing majority of prenatal opioid exposure cases^18,19^. Initiating opioid exposure at later stages of pregnancy overlooks the effect of opioids on earlier embryonic developmental processes. These weaknesses underlie the call for improved animal models of prenatal opioid exposure, which encompass all stages of prenatal brain development^11^.

We developed a more translational mouse model that resembles the typical pattern of opioid use in a pregnant woman who is first dependent on oxycodone, then begins methadone maintenance treatment, and subsequently becomes pregnant while maintained on methadone. We found that PME reduces physical growth in offspring which persists into adolescence and disrupts the development of locomotor activity and ultrasonic vocalization (USV) during the preweaning period. Furthermore, PME delays the development of specific sensorimotor milestones. These impairments were concurrently associated with alterations in layer V motor cortical neuron intrinsic properties and local connectivity.

## Results

### Impact of PME on Pregnancy and Litter Characteristics

An overview of model development and workflow representing the total animals used for these studies is found in **Fig. 1. Table 1** summarizes pregnancy and litter characteristics. Opioid treatment did not reduce pregnancy rates (chi square test: χ^2^_(1, n=95)_=0.904, p=0.34) or instances of obstructed labor (chi square test: χ^2^_(1, n=70)_ =2.14, p=0.14). Litter sizes at birth (unpaired t test: t_61_=1.36, p=0.18) and total neonatal deaths (chi square test: χ^2^_(1, n=406)_=0.201, p=0.65) were not affected by opioid treatment. There was a trend for fewer PME males surviving to P7 (chi square test: χ^2^_(1, n=336)_=2.71, p=0.10). However, the proportion of sexes at birth between treatment groups was not significantly different (chi square test: χ^2^_(1, n=284)_=0.010, p=0.92) suggesting the reduction of male offspring at P7 may be attributed to the greater postnatal deaths in males relative to females in PME litters (chi square test: χ^2^_(1, n=37)_=4.79, p=0.03).

**Figure 1.**
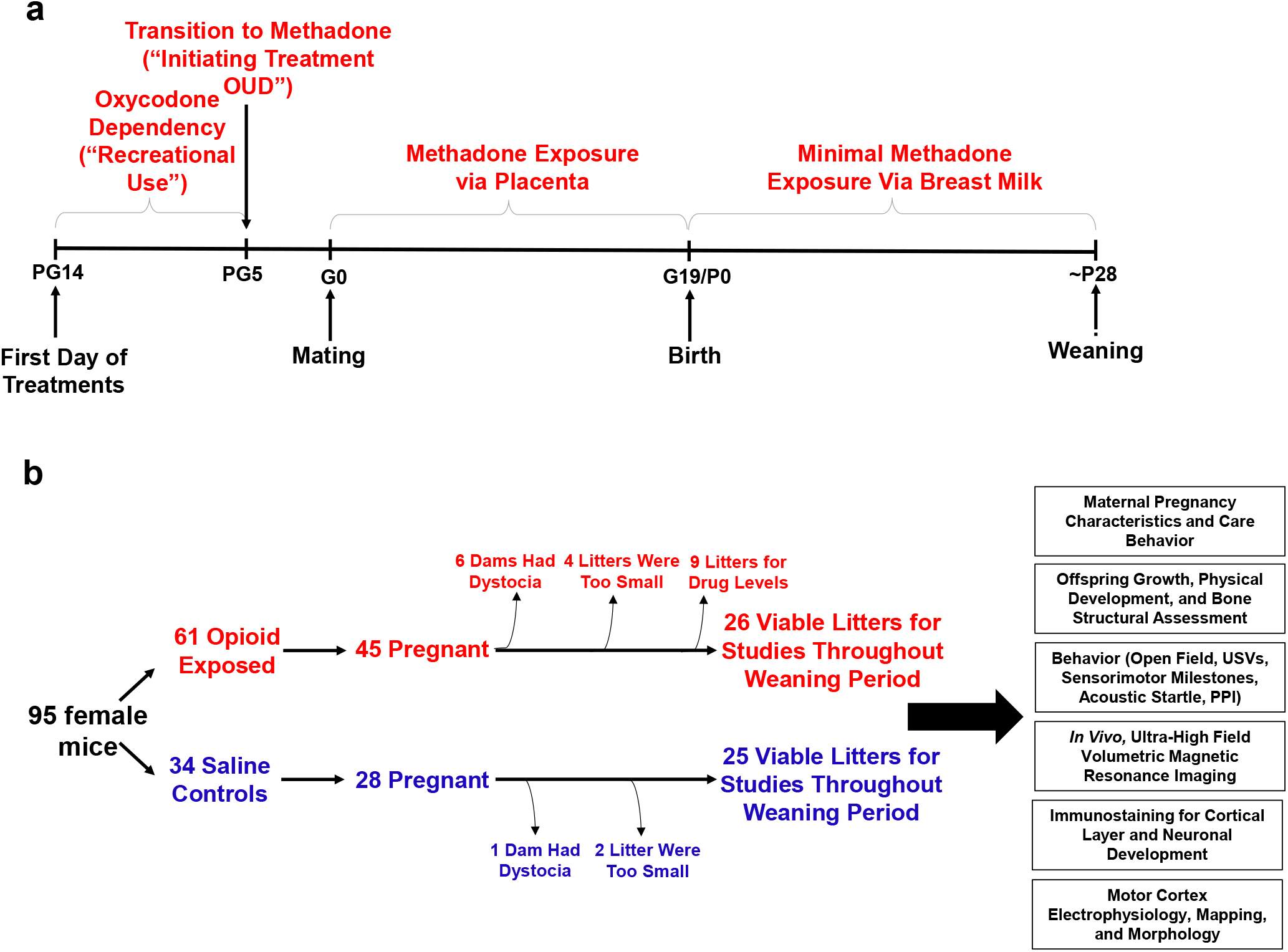
Study Overview. **a** Timeline briefly describing the generation of mice with prenatal methadone exposure (PME). Approximately 15 days prior to mating (pregestational day 14), female mice begin treatment with oxycodone to model recreational opioid use. Following nine days of oxycodone, mice began methadone to simulate treatment of opioid use disorder (OUD). Following five days of methadone administration, the females were mated (gestational day 0) and treatment continued throughout gestation passively exposing the developing embryo and fetus to methadone. Following birth, dam methadone treatment continued which provided minimal, but measurable methadone exposure to offspring (see **Fig 2.** and **Supplementary Table 1**). Offspring were weaned shortly at approximately postnatal day 28. All doses were given subcutaneously twice daily. Control animals underwent an identical timeline with the exception of receiving saline injections. **b** Flow chart reviewing studies completed in the present study using offspring generated from the timeline in **Fig. 1a**. Female mice were randomly assigned to either opioid (oxycodone → methadone) or saline treatment. Of the 45 opioid treated mice which became pregnant, six were removed from the study due to obstructed labor (dystocia), four litters were too small (<3 pups), and nine were chosen at random to assess methadone tissue and plasma concentrations. Of the 28 pregnant control mice, one experienced dystocia and two litters were too small. The offspring from the remaining litters (25-26 per exposure) were pseudo-randomly allocated to the behavioral and biological studies indicated in the black boxes (≥4 litters per exposure with equal representation of both sexes were used for each experiment). See **Table 1** for detailed description of litter characteristics. *PG*, pregestation; *G*, gestation; *P*, postnatal, *OUD*, opioid use disorder; *USVs*, ultrasonic vocalizations; *PPI*, prepulse inhibition.

**Table 1.** Pregnancy and Litter Characteristics. All data was analyzed using Chi-Square tests except litter size which was analyzed using an unpaired, independent t test (value represents mean ± SEM). Sex was not determined in earlier cohorts of animals until nipple marks began to appear (~P7). However, given the potential sex differences in survival, for later cohorts we began determining sex at P0 and P7 which led to a lower total sample size of litters at the P0 date of sex determination. Postnatal deaths occur within approximately 72 hours after birth. As not all litters survived to ~P7 when sex was determined in offspring, the total sample size of litters from which postnatal deaths were examined is larger than the total sample size of litters for P7 sexes.

|  | PME | PSE | p-value |
| --- | --- | --- | --- |
| **Total Pregnancies (%)** | 45/61 mated (73.8%) | 28/34 mated (82.4%) | p=0.34 |
| **Obstructed Labor (%)** | 6/42 pregnancies (14.3%) | 1/27 pregnancies (3.6%) | p=0.14 |
| **Litter Size** | 7.50 ± 0.34 (n=36 litters) | 6.85 ± 0.31 (n=27 litters) | p=0.18 |
| **P0 Male:Female Ratio (% Male)** | 96M:66F from 21 litters (59.3%) | 73M:49F from 18 litters (59.8%) | p=0.58 |
| **P7 Male:Female Ratio (% Male)** | 90M:90F from 26 litters (50.0%) | 92M:64F from 25 litters (59.0%) | p=0.10 |
| **Postnatal Deaths (%)** | 50/248 from 33 litters (20.2%) | 29/158 from 27 litters (18.4%) | p=0.65 |
| **Male:Female Postnatal Death Ratio (% Male Deaths)** | 18M:5F from 21 litters (78.3%) | 6M:8F from 18 litters (42.9%) | **p=0.03** |

### Methadone and EDDP Concentrations in Dams and Offspring

Methadone and 2-ethylidene-1,5-dimethyl-3,3-diphenylpyrrolidine (EDDP, main metabolite of methadone) concentrations in the placenta, plasma, and brain for dams and offspring on G18, P1, and P7 are presented in **Supplementary Table 1** and **Fig. 2a**. Maternal brain and plasma levels were highest at G18 (248.9 ± 54.1 ng/g and 63.5 ± 11.3 ng/mL, respectively) and lower slightly after giving birth, remaining relatively stable at P1 and P7. Methadone accumulated in the brain relative to the plasma for dams at all timepoints (**Fig. 2a**). The placenta significantly retained methadone (3862.1 ± 258.4 ng/g) and EDDP (1124.0 ± 84.4 ng/g; **Fig. 2a,b**). The fetal brain methadone concentration was 2100.8 ± 237.6 ng/g which was substantially higher than the maternal brain on G18 (**Fig 2a**). Offspring brain levels dropped precipitously to 7.9 ± 0.6 ng/g and 3.1 ± 0.3 ng/g on P1 and P7, respectively. Although we were not able to collect a sufficient plasma volume on G18, minimal methadone plasma levels in offspring were quantified postnatally. Placental levels were predictive of fetal brain methadone levels at G18 (R^2^=0.39, p<0. 0078; **Fig. 2c**). Offspring plasma methadone was predictive of offspring brain methadone at P1 (R^2^=0.45, p<0.0084; **Fig. 2d**), but not on P7 (R^2^=0.008, p=0.74; **Fig. 2e**).

**Figure 2.**
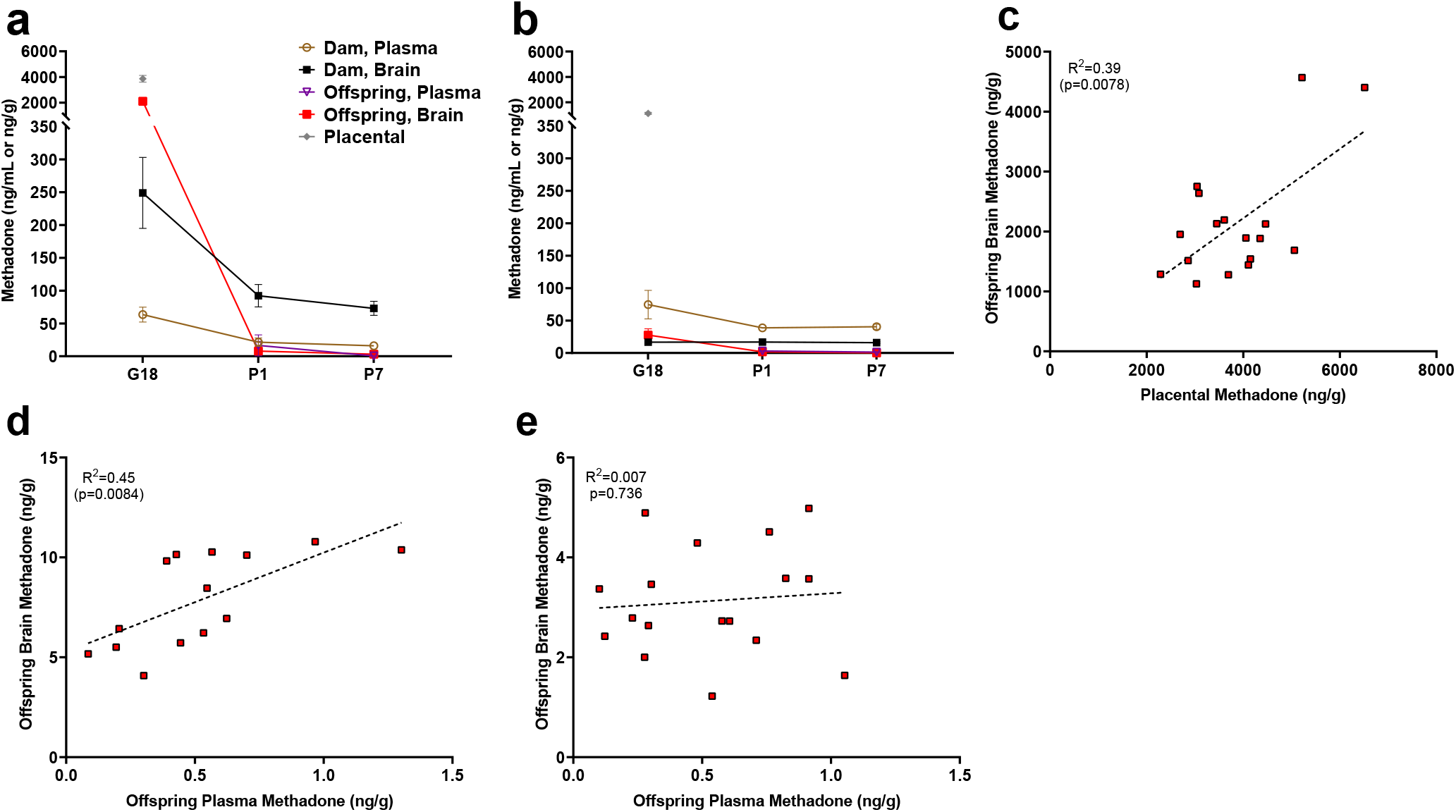
Relationship Between Placental, Plasma, and Brain Methadone and Metabolite Levels in Dams and Offspring During Gestation and the Postnatal Period. **a** Methadone and **b** EDDP (2-ethylidene-1,5-dimethyl-3,3-diphenylpyrrolidine, main metabolite of methadone) concentrations in the plasma, brain, and placenta of dams and offspring on G18 (approximately 1 day before birth), P1 (approximately 1 day after birth), and P7. Methadone highly accumulated in the fetal compartment relative to dam concentrations, but methadone concentrations dropped precipitously following birth. EDDP accumulated in the placenta. Data points indicate mean ± SEM. **c** Placental methadone concentrations predicted fetal brain concentrations on G18 (R^2^=0.39, p=0.0078). **d,e** Offspring plasma methadone concentrations predicted offspring brain methadone on P1 (R^2^=0.45, p=0.0084), but not on P7 (R^2^=0.007, p=0.736). All tissue and blood samples were collected 2.5 hours following the morning administration of methadone. (n=3 dams + their respective litters per timepoint; n=17-20 offspring samples at G18, n=15 offspring at P1, n=17-18 offspring at P7). Data are collapsed across offspring sex. The limit of quantification for methadone and EDDP detection was 0.1 ng/mL and 0.05 ng/mL in the plasma, respectively, and 0.08 ng/sample and 0.04 ng/sample of placenta and brain for both methadone and EDDP.

### Maternal Characteristics in Opioid-Treated Dams

The opioid treatment strategy did not significantly impact maternal weights during the course of the study (rmANOVA: Treatment, F_(1,42)_=0.00117, p=0.97; Time, F_(2.92,122.4)_=266.1, p<0.0001; Interaction, F_(22,924)_=0.510, p=0.97; **Fig. 3a**). Measures of food consumption revealed a significant effect of opioid treatment (rmANOVA: Treatment, F_(1,33)_=5.09, p=0.03; Time, F_(7,231)_=413, p<0.0001; Interaction, F_(7,231)_=2.48, p=0.02; **Fig. 3b**) with an increased food consumption during the postnatal period in opioid-treated dams reaching significance on postnatal week 2 (Sidak’s post-hoc test, p=0.004). Maternal care-related behavior on P3 was assessed to examine the potential impact of methadone on offspring care. There were no apparent differences in nest quality (Mann-Whitney test: U=53, p=0.15; **Fig. 3c**) or average latency to retrieve pups removed from the nest (Mann-Whitney test: U=77, p=0.97; **Fig. 3d**). No placentas remained in the cages 24 hours after birth in either group indicating normal placentophagy. Oxycodone treatment prior to gestation (PG14-PG6; see **Fig. 1a**) induced opioid dependency in dams as demonstrated by the significantly increased withdrawal behaviors following naloxone treatment (**Fig. 3e**). Offspring from mothers that were assessed for oxycodone dependency were not used in subsequent analyses to avoid any impact that pregestational opioid withdrawal could have on offspring outcomes. Methadone maintained opioid dependency as evidenced by the significantly increased naloxone-precipitated withdrawal behaviors observed in opioid-treated females at post-weaning (ANOVA: Treatment, F_(1,26)_=38.1, p<0.0001; Stage, F_(1,26)_=0.361, p=0.55; Interaction, F_(1,26)_=0.337, p=0.56; **Fig. 3e**).

**Figure 3.**
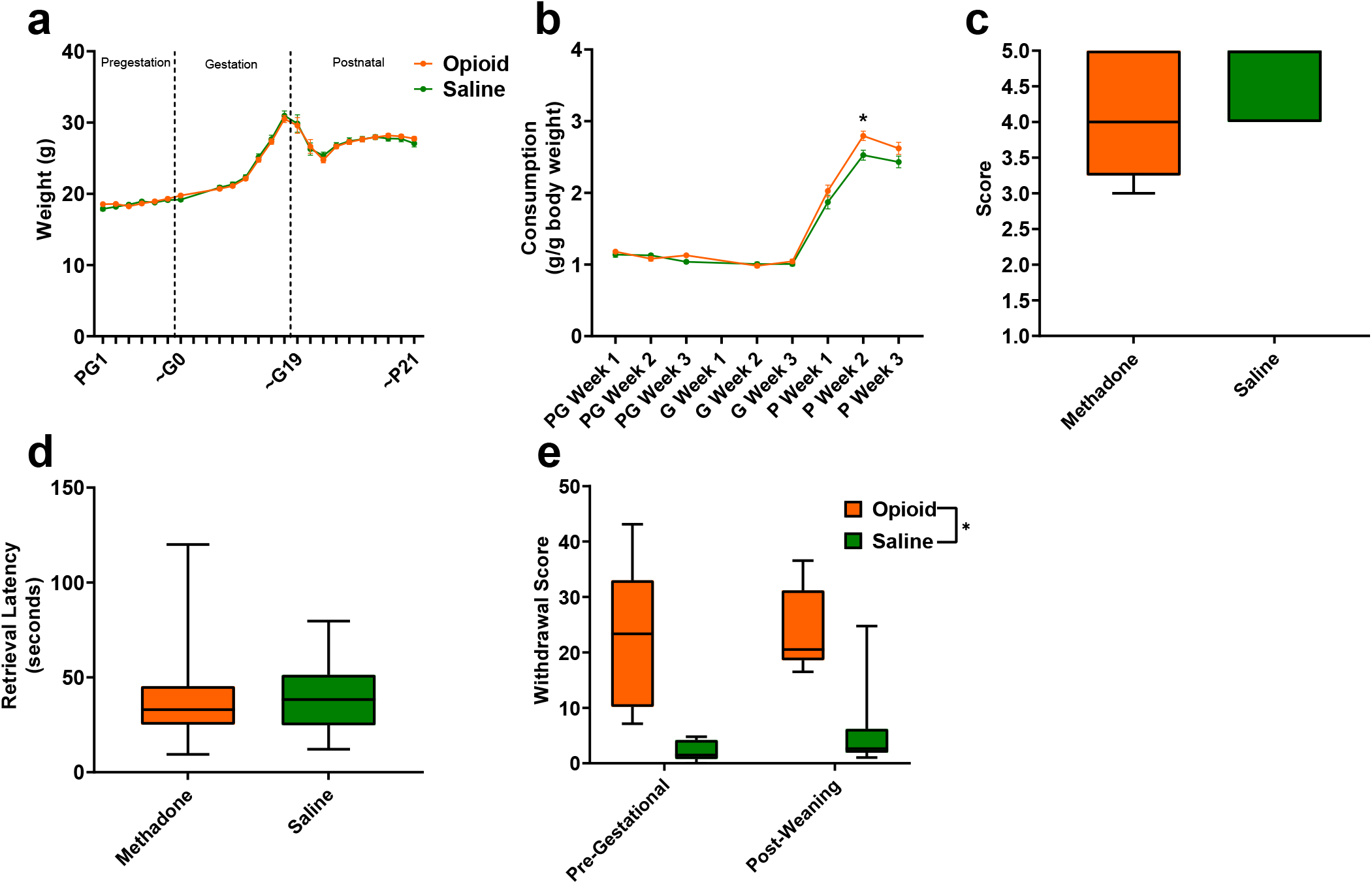
Opioid Treatment Induced Dependency and Reduced Gestational Weight Gain but Did Not Alter Maternal Care. **a** Measures of maternal weights over the course of the study revealed no significant effects of opioid treatment (n=23 opioid, 21 saline mice); however, **b** Maternal food consumption was not significantly affected during pregestation and gestation but was significantly increased in opioid-treated dams following birth (rmANOVA: Interaction, p=0.02; Sidak’s post hoc: P week 2, p=0.004, n=19 opioid, 16 saline mice). **c,d** Opioid-treated dams demonstrated no differences in **c** nest quality or **d** latency to retrieve pups removed from the nest on P3 suggesting maternal care behavior was grossly intact (n=12 opioid, 13 saline mice). **e** The nine days of pregestational oxycodone treatment induced maternal opioid dependency and the oxycodone treatment plus the transition to methadone throughout gestation and the pre-weaning period was sufficient to maintain opioid dependence as determined by measures of naloxone-precipitated withdrawal behaviors (ANOVA: Treatment, p<0.0001, n=7-8 opioid, 7-8 saline mice). *p<0.05. **a,b** Data points indicate mean ±SEM. **c-e** box plots indicate 25th to 75th percentiles with whisker characterizing the minimum and maximum value.

### Physical Development of Offspring

The clinical evidence for the effect of prenatal opioid exposure on fetal growth has revealed mixed findings^20^; therefore, we first examined the impact of PME on offspring physical development. As there were no significant effects of sex, the data on weight and lengths during the pre-weaning period were pooled. PME significantly reduced weights and reduced weight gain during the pre-weaning period (rmANOVA: Exposure, F_(1,111)_=3.96, p=0.049; Time, F_(1.344,149.2)_=3177, p<0.0001; Interaction, F_(6,666)_=2.89, p=0.0087; **Fig. 4a**). The reduced body weight in PME offspring persisted into adolescence as weights remained consistently lower across sexes at P35 and P49 (rmANOVA: Exposure, F_(1,50)_=7.52, p=0.0085; Sex, F_(1,50)_=131, p<0.0001, Time, F_(1,50)_=673, p<0.0001; Time x Sex, F_(1,50)_=62.3, p<0.0001; Time x Exposure, F_(1,50)_=1.60, p=0.21; Exposure x Sex, F_(1,50)_=0.00956, p=0.92; Time x Exposure x Sex, F_(1,50)_=0.20, p=0.66; **Fig. 4a**). Similarly, body length was reduced by PME (rmANOVA: Exposure, F_(1,111)_=12.24, p=0.0007; Time, F_(2.676,297.0)_=4719, p<0.0001; Interaction, F_(6,666)_=0.987, p=0.433; **Fig. 4b**).

As chronic opioid use is associated with bone pathology^21^, we collected femurs from P7 and P35 offspring to assess bone structure and density. Although femur length was not impacted by PME (unpaired t test: t_23_=1.126, p=0.27; **Supplementary Fig. 1a**), whole bone volume tended to be lower in P7 PME offspring (unpaired t test: t23=1.89, p=0.072; **Supplementary Fig. 1b)**. By P35, structural bone measures were similar between exposure groups including distal femur metaphysis bone volume (ANOVA: Exposure, F_(1,16)_=1.94, p=0.18; Sex, F_(1,16)_=13.4, p=0.0021; Interaction, F_(1,16)_=2.17, p=0.16; **Supplementary Fig. 1c**), trabecular bone volume (ANOVA: Exposure, F_(1,16)_=1.30, p=0.27; Sex, F_(1,16)_=25.7, p=0.0001; Interaction, F_(1,16)_=0.126, p=0.73; **Supplementary Fig. 1d**), cortical bone area (ANOVA: Exposure, F_(1,16)_=0.000, p=0.99; Sex, F_(1,16)_=13.4, p=0.0021; Interaction, F_(1,16)_=2.04, p=0.17; **Supplementary Fig. 1e**), and cortical thickness (ANOVA: Exposure, F_(1,16)_=0.0435, p=0.84; Sex, F_(1,16)_=19.0, p=0.0005; Interaction, F_(1,16)_=0.790, p=0.39; **Supplementary Fig. 1f**).

A recent report suggested opioid-exposed infants have a higher prevalence of orofacial clefting which may reflect an effect of prenatal drug exposure on craniofacial development^22^. However, we did not observe any effect on the day both eyes opened (unpaired t test: t_49_=1.67, p=0.10), both ears moved to their final erect position and external auditory canals were patent (unpaired t test: t_49_=0.910, p=0.37), or when bottom incisor teeth erupted (unpaired t test: t_49_=1.05, p=0.30; **Supplementary Fig. 2**) indicating measures of mouse postnatal craniofacial development progress normally in our PME model.

**Figure 4.**
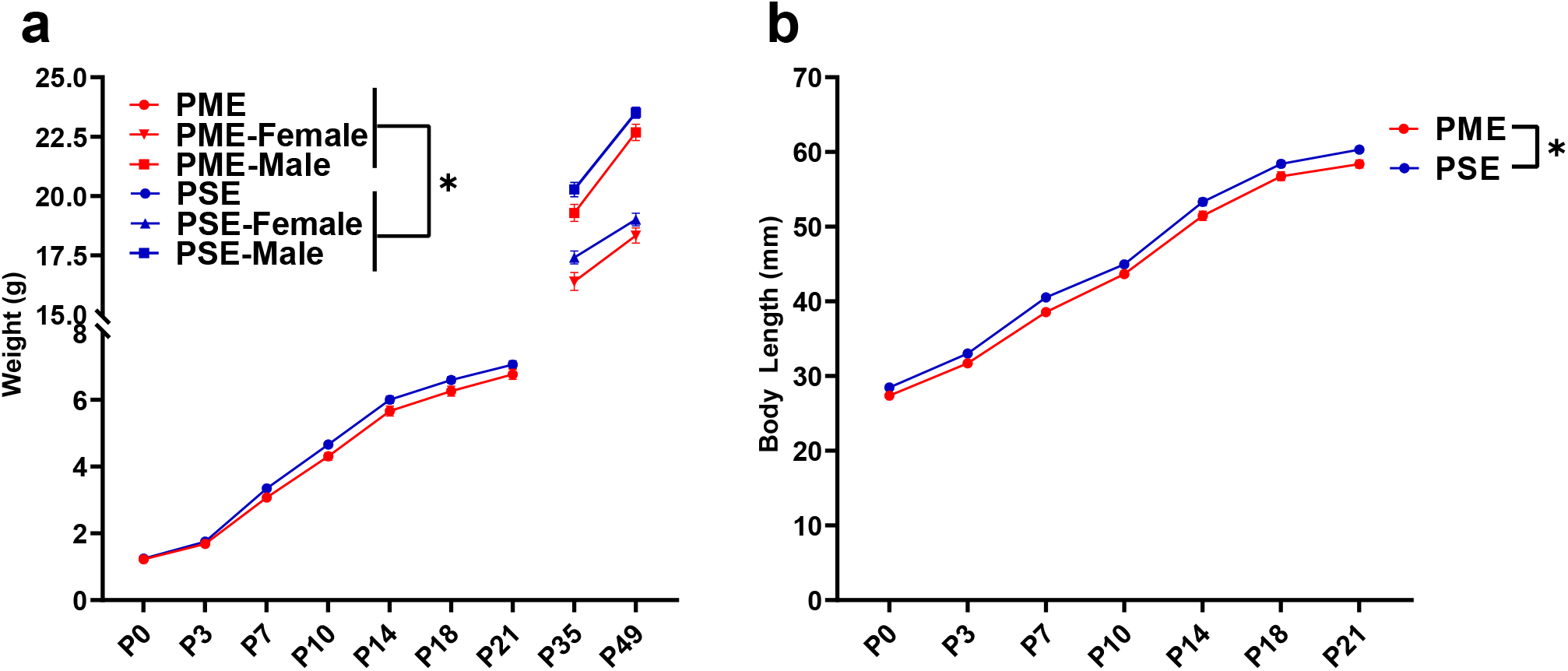
PME Produced a Persistent Reduction in Offspring Weight and Body Length. **a** PME reduced offspring weight during the preweaning period (rmANOVA: Exposure, p=0.049; Interaction, p=0.0087, n=57 PME (25M:32F), 56 PSE (29M:27F)) and this effect on weight persisted into adolescence in both sexes (rmANOVA: Exposure, p=0.0085, n=30 PME (16M:14F), 24 PSE (15M:9F)). **b** PME reduced offspring body length during the preweaning period (rmANOVA: Exposure, p=0.0007, n=57 PME (25M:32F), 56 PSE (29M:27F)). *main effect of exposure p<0.05 Data points indicate mean ±SEM.

### Behavioral Development in Offspring

As methadone levels rapidly dropped from G18 to P1 in offspring, we sought to determine if PME offspring demonstrated behaviors consistent with NOWS such as hyperthermia or myoclonic jerks (twitching or jerking of the limbs or whole-body) which are indicators of NOWS in humans^23^. PME offspring displayed a relative hyperthermia at baseline that was maintained after two minutes of isolation (rmANOVA: Exposure, F_(1,48)_=34.1, p<0.0001; Time, F_(1,48)_=577, p<0.0001; Interaction, F_(1,48)_=0.127, p=0.723; **Fig. 5a**). A higher number of twitches/jerks were observed in P1 PME offspring (unpaired t test: t_49_=2.04, p=0.047; **Fig. 5b**) reminiscent of the myoclonic jerks which are common among human neonates experiencing NOWS from methadone^23^. In association with a rapid drop in brain methadone levels from G18 to P1, the relative hyperthermia and increased twitches/jerks suggests PME offspring may experience opioid withdrawal at P1.

Preclinical studies have indicated locomotor activity may be different in prenatal opioid-exposed offspring^24^, however, no studies to date have examined the trajectory of locomotor development during early life in opioid-exposed offspring. Using an open field, we measured the development of motor activity by repeatedly testing animals on P1, P7, P14, and P21. Locomotor activity was significantly altered during the pre-weaning period in PME offspring (rmANOVA: Exposure, F<1,4β)=5.18, p=0.027; Time, F_(1.38,67.84)_=299, p<0.0001; Interaction, F_(3,147)_=10.5, p<0.0001) with PME offspring demonstrating significantly reduced activity at P1 but greater activity at P7 and P21 (Sidak’s post hoc test: p=0.048, p=0.037, p=0.009, respectively; **Fig. 5c**). Inspection of the P1 videos indicated the reduction in motor activity was likely a result of poor coordination as PME displayed a greater propensity to fall over earlier in the testing session. Although 88% of PME and 80% of PSE offspring fell over by the end of five-minute session on P1 (chi square test: χ2_(1, n=51)_=0.690, p=0.41; **Supplementary Fig. 3**), 53.5% of PME compared to 28% of PSE animals had fallen over immediately upon placement into the arena (chi square test: χ^2^ (1, n=51)=0.3.52, p=0.061; **Supplementary Fig. 3b**). There was a nonsignificant reduction in the latency to fall over in PME offspring (Mann-Whitney test: U=248.5, p=0.136; **Supplementary Fig. 3c**).

In addition to the hyperactivity on P21, PME offspring exhibited significantly more vertical jumps at P21 (Mann-Whitney test: U=53, p<0.0001; **Fig. 5d**) further indicating a hyperactive phenotype is present in juvenile PME offspring similar to what has been reported in opioid-exposed children^25,26^. These jumps may represent escape attempts as some animals were able to jump and reach the top of the arena walls. When this occurred, the animal was outside of the frame of video and our tracking software was not able to record activity. Therefore, the total distance traveled at P21 in the PME animals may actually be greater than we are able to report. As increased anxiety-like behavior has also been reported in some prenatal opioid exposure models^18,24^, we further examined open field videos for thigmotaxis, grooming, and rearing behaviors. Thigmotaxis was affected by PME (rmANOVA: Exposure, F_(1,49)_=0.17.33, p=0.0001; Time, F_(1.41,68.9)_=236.3, p<0.0001; Interaction, F_(2,98)_=12.01, p<0.0001), with PME animals spending significantly more time at P7 near the arena walls than PSE animals (Sidak’s post hoc test: p=0.0004 **Supplementary Fig. 4**) However, this likely reflects the relative hypoactivity of PSE animals which demonstrated very little activity on P7 and mostly remained in the arena center where offspring were placed at the beginning of each trial. Both grooming (Mann-Whitney test: U=173.5, p=0.0037) and unsupported rearing (unpaired t test: t_49_=2.48, p=0.017), but not supported rearing (unpaired t test: t_49_=0.352, p=0.73) were reduced in PME mice which likely reflects an increase in time spent in active locomotion and jumping instead of grooming and rearing in PME mice as opposed to anxiolysis (**Supplementary Fig. 4b-d**).

When removed from the dam, pups emit separation-induced USVs which signal distress and encourage retrieving behavior from the mother. USVs typically peak around P8 and extinguish by the third week of life^27^. The production of USVs in the open field was also significantly altered by PME (rmANOVA: Exposure, F_(1,47)_=0.380, p=0.54; Time, F_(3,141)_=13.3, p<0.0001; Sex, F_(1,47)_=4.24, p=0.045; Time x Exposure, F_(3,141)_=3.01, p=0.032; Time x Sex, F_(3,141)_=0.636, p=0.59; Exposure x Sex, F_(1,47)_=1.57, p=0.22; Time x Exposure x Sex, F_(3,141)_=0.947, p=0.42; **Supplementary Fig. 5** and collapsed on sex in **Fig. 5e**). Female PME offspring vocalized significantly more than both male and female PSE offspring at P7 (Sidak’s post hoc test; p=0.0045 & p=0.026, respectively) but this difference was not observed in male PME offspring relative to either sex of PSE offspring (Sidak’s post hoc test; both p>0.999, **Supplementary Fig. 5**). When collapsed on sex, vocalizations are nonsignificantly reduced at P1 and significantly increased at P7 in PME offspring (Sidak’s post hoc test: p=0.079 & p=0.015, respectively; **Fig. 5e**). Although female PME offspring do not produce significantly more USVs than male PME offspring at P7 (Sidak’s post hoc: p=0.60), the sex-collapsed effect described on P7 is likely driven by the high rate of USV production in female PME offspring (**Supplementary Fig. 5**). We next performed further analysis on P7 USVs as this day differed between exposure groups. Total activity correlated significantly with total calls on P7 across all animals (Pearson’s r=0.41, p=0.0028; **Supplementary Fig. 6a**) suggesting that there may be a relationship between hyperactivity and hyper-vocalizations among PME offspring. USV call types were classified into categories based on frequencies, to cluster the types of USVs emitted on P7^28^. The call classification analysis revealed the increase in USVs on P7 may be a result of a higher number of complex, flat, step down, and upward ramp calls emitted by PME offspring (unpaired t tests: t_49_=2.54, p=0.014; t_49_=2.98, p=0.0045; t_49_=2.74, p=0.0086; t_49_=2.09, p=0.041; **Supplementary Fig. 6b**). Markov chains of call type transitions (‘syntax’) was further analyzed on P7 which also revealed differences in the transition between call types between exposure groups (**Supplementary Fig. 6c**).

Beginning on P3 and through P14, offspring began a battery of daily developmental tests assessing the development of sensorimotor milestones. Offspring with PME demonstrated delayed acquisition of surface righting (unpaired t test: t48=4.69, p<0.0001), forelimb grasp (Mann-Whitney test: U=166, p=0.0029), and cliff aversion (Mann-Whitney test: U=161, p=0.0026; **Fig. 5f**). No differences in the acquisition of the extinguishment of pivoting (Mann-Whitney test: U=238, p=0.14) or negative geotaxis behavior (Mann-Whitney test: U=270.5, p=0.42) were observed. As basic locomotor activity is not impaired in PME offspring, these data indicate that PME offspring lack the ability to produce more complex, coordinated motor behaviors in response to multimodal sensory input.

A recent investigation of prenatal THC exposure revealed deficits in sensorimotor gating^29^; therefore, as sensorimotor performance was impaired in PME offspring, we subsequently tested prepulse inhibition (PPI), a sensorimotor gating task. In a cohort of P28-P29 offspring, we found that PME did not alter the acoustic startle response (rmANOVA: Exposure, F_(1,36)_=0.129, p=0.72; Intensity, F_(1.830,65.87)_=185.5, p<0.0001; Interaction, F_(6,216)_=0.147, p=0.99; **Supplementary Fig. 7a**). No exposure-related effects were found on the percent PPI suggesting sensorimotor gating is intact in this model (rmANOVA: Exposure, F_(1,36)_=0.282, p=0.60; Prepulse Intensity, F_(1.860,66.97)_=30.8, p<0.0001; Interaction, F_(2,72)_=0.770, p=0.47; **Supplementary Fig. 7b**).

**Figure 5.**
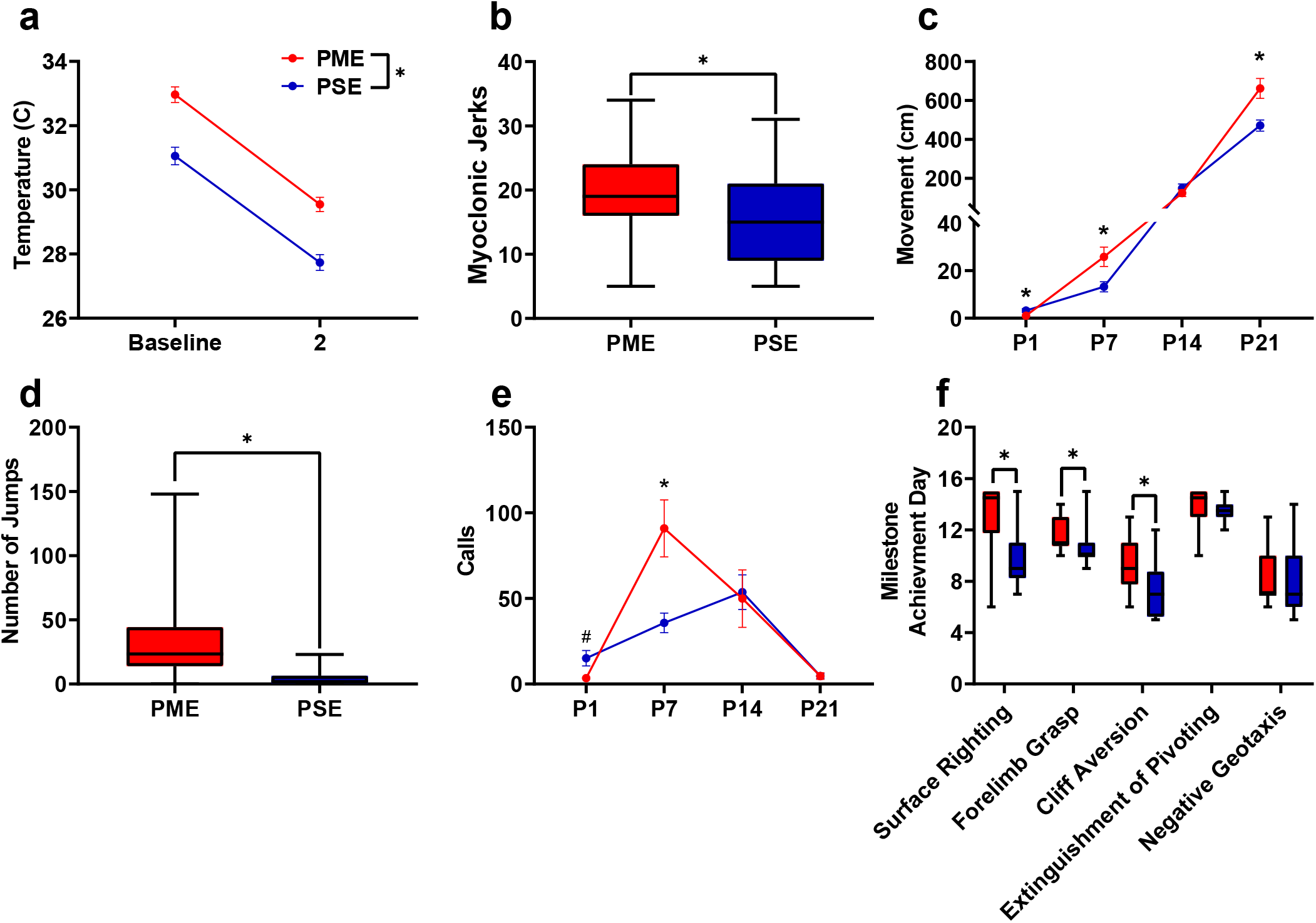
PME Disrupted Behavioral Development in Offspring. **a** Offspring showed significantly higher surface body temperature when removed from the nest (baseline) and continued to maintain a higher surface body temperature following two minutes of isolation at P1 (rmANOVA: Exposure, p<0.0001, n=25 (7M:18F) PME, 25 PSE (15M:10F) mice). **b** PME offspring show a greater number of twitches/jerks at P1 similar to the myoclonic jerks reportedly observed in human neonates experiencing opioid withdrawal (unpaired t test: p=0.047, n=26 (7M:19F) PME, 25 PSE (15M:10F) mice). **c** Offspring were repeatedly tested in a modified open field during the first three weeks of life to examine the development of locomotor activity and ultrasonic vocalization (USV) production. **c** PME altered the development of locomotor activity (rmANOVA: Exposure, p=0.027; Interaction, p<0.0001) with PME offspring showing reduced activity at P1 but significantly greater activity at P7 and P21 (Sidak’s post hoc test: p=0.048, p=0.037, p=0.009, respectively, n=26 (7M:19F) PME, 25 PSE (15M:10F) mice). **d** Subsequent manual scoring of videos on P21 revealed significantly greater number of vertical jumps during the five-minute session in PME mice further supporting a hyperactive phenotype at P21 (Mann-Whitney test: p<0.0001, n=26 (7M:19F) PME, 25 PSE (15M:10F) mice). **e** PME offspring showed differences in the production of separation-induced USV calls (rmANOVA: Interaction, p=0.032) with less total calls emitted on P1 and significantly more calls emitted on P7 (Sidak’s post hoc test: p=0.079, p=0.015, n=26 (7M:19F) PME, 25 PSE (15M:10F) mice). **f**. Acquisition of the surface righting (unpaired t test: p<0.0001), forelimb grasp, and cliff aversion (Mann-Whitney test: p=0.0029 & p=0.0026, respectively) were significantly delayed in PME offspring (n=26 (7M:19F) PME, 24 PSE (14M:10F) mice). * p<0.05, # p=0.079. Data points indicate mean ±SEM. Box plots indicate 25th to 75th percentiles with whisker characterizing the minimum and maximum value.

### Brain Anatomical Development in Offspring

Numerous brain regions and neural mechanisms could contribute to the altered activity and sensorimotor development described in PME offspring; therefore, we began our investigation by probing structural differences in grey matter regions across the brain using in vivo, ultra-high field volumetric MRI. Volumes of interest (VOIs; See **Supplementary Fig. 8** for segmentations) were used to determine if differences in various grey matter regions were present in offspring at P28-P30. No significant differences in VOIs across any of the brain regions examined were discovered suggesting gross grey matter structure is mostly unaffected by PME (unpaired t tests: see **Supplementary Table 2** for full test statistics; **Supplementary Fig. 9**).

To determine whether PME could alter cortical development, we examined cortical laminations and neuron densities in distinctive cortical layers of P22-P24 offspring with triple immunostaining of NeuN (neuronal marker), Cux1 (a cortical layer II-IV marker), and Draq5 (nuclei stain). We focused on the anterior cingulate cortex (ACC), primary somatosensory cortex (S1), and primary motor cortex (M1) for their involvement in sensorimotor behavior^30-32^ (**Fig. 6, Supplementary Fig. 10-11)**. In ACC, both the distribution patterns of Cux1 (rmANOVA: Exposure, F_(1,250)_=0.000, p=0.99; Bin, F_(9,250)_=369 p<0.0001; Interaction, F_(9,250)_=1.73, p=0.082) and NeuN densities (rmANOVA: Exposure, F_(1,250)_=0.0786, p=0.78; Bin, F_(9,250)_=104 p<0.0001; Interaction, F_(9,250)_=1.77, p=0.075) are similar between PME and PSE offspring. The cortical lamination and neuron densities in S1 of PME are also similar to PSE (NeuN distribution-rmANOVA: Exposure, F_(1,240)_=0.389, p=0.53; Bin, F_(9,240)_=55.8, p<0.0001; Interaction, F_(9,240)_=1.58, p=0.12; Cux1 distribution-rmANOVA: Exposure, F_(1,240)_=0.000, p>0.99; Bin, F_(9,240)_=324 p<0.0001; Interaction, F_(9,240)_=1.44, p=0.17). In M1, we found a significant reduction in NeuN densities in cortical layer II-IV (unpaired t test: t_26_=2.519, p=0.018; **Fig. 6c**) but not layer V (unpaired t test: t_26_=0.734, p=0.47; **Fig. 6d**). There is also no difference in the distributions of Cux1^+^ cells (unpaired t test: t_26_=0.459, p=0.65; **Fig. 6f**). These data suggest that cortical lamination develop normally in PME for ACC, S1, and M1 regions. However, the neuronal density is reduced in the upper cortical layer in PME M1.

**Figure 6.**
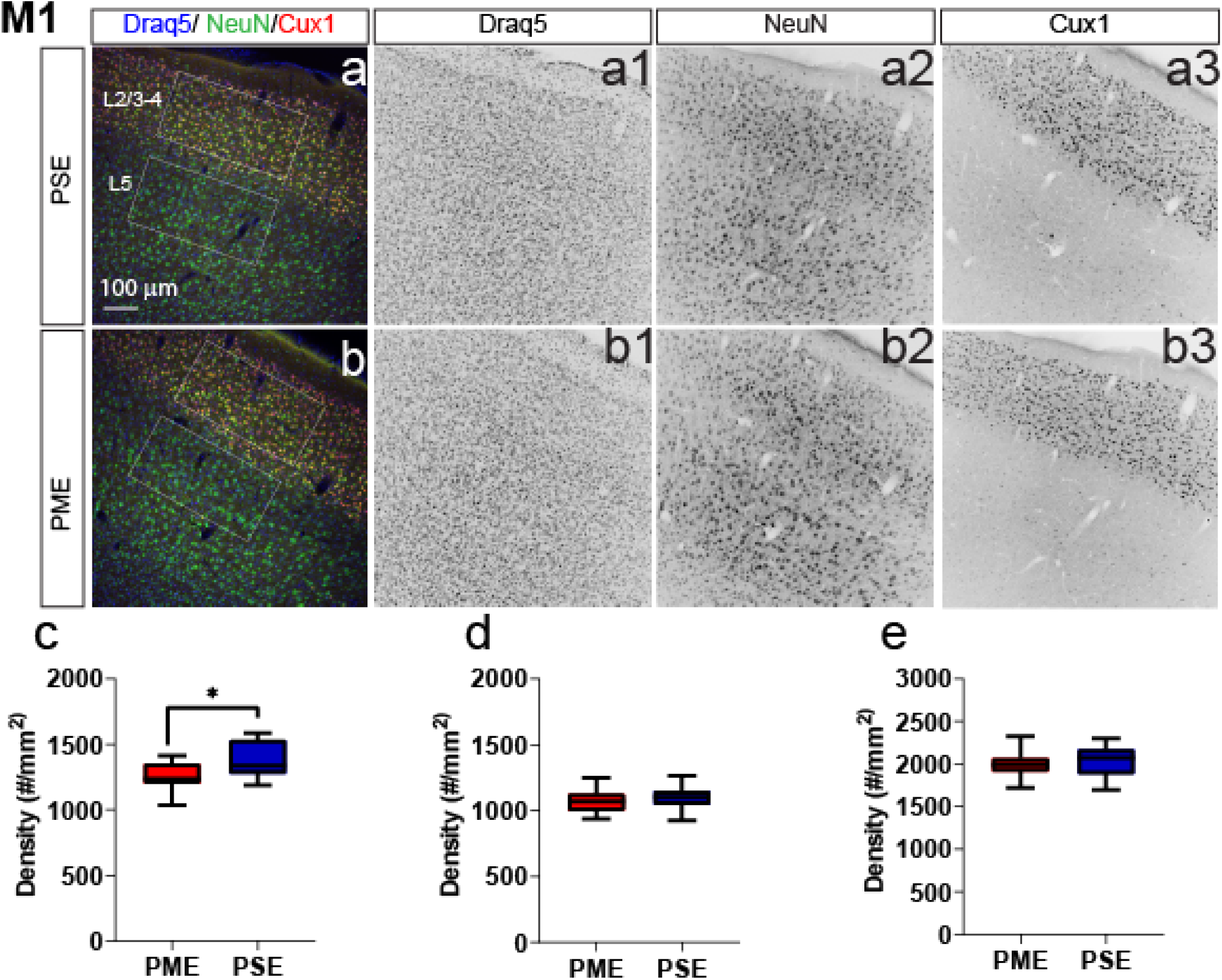
Neuronal Density is Reduced in the Primary Motor Cortex (M1). Representative slices of the M1 in **a** PSE and **b** PME offspring demonstrating Draq5 (blue, a1,b1: marker of nuclei), NeuN (green, a2,b2,: marker of post-mitotic neurons), and Cux1 (red, a3,b3: marker of upper cortical layers, typically layer 2/3). White boxes represent the areas used for quantification. **c** A significant decrease in NeuN-positive cells was found in layer 2/3 of PME offspring (unpaired t test: p=0.018). No differences in density of **d** NeuN in layer 5 or **e** Cux1 in layer 2/3 were found (n=7 (2M:5F) PME, 7 PSE (3M:4F). Box plots indicate 25th to 75th percentiles with whisker characterizing the minimum and maximum value.

### Motor Cortex Neuronal Intrinsic Properties and Local Circuitry

Given the significant alterations in sensorimotor behavior in offspring with PME, we next examined intrinsic properties, local circuitry, and morphology of layer 5B thick-tufted pyramidal neurons of the motor cortex in P21-P26 offspring using whole-cell patch clamp electrophysiology and laser scanning photostimulation for optical mapping of local circuitry (See **Fig. 7a,b** for schematic diagram, and **Supplementary Fig. 10** for neuron morphological reconstructions used for Sholl analyses). Thick-tufted layer 5B neurons indicate neurons with specific subcortical projection targets^33-35^, and most likely represented M1 pyramidal tract corticospinal neurons^36^. Representative current-clamp traces from whole-cell recordings of action potential firing and sub-threshold voltage responses are shown in **Supplementary Fig. 13a & Fig. 7c,** respectively. Recording analyses showed that L5 M1 neurons from PME mice displayed significantly reduced firing rates (number of APs) in response to injected current compared to PSE offspring (rmANOVA: Exposure, F_(1, 66)_=0.707, p=0.40; Current, F_(2.198, 145.1)_=1109 p<0.0001; Interaction, F_(12, 792)_=1.78, p=0.047, **Supplementary Fig. 13b**). These findings suggest PME neurons are characterized by reduced intrinsic excitability at higher injected currents; however, no posthoc tests reached the level of significance. PME neurons showed exhibited significantly reduced input resistance compared to PSE cells (**Fig 7d**, see **Supplementary Table 4** for stats), indicating that current was translated into a smaller change in voltage across the membrane of PME neurons. Additionally, subthreshold responses of L5 M1 neurons showed prominent “sag” and “overshoot” of the membrane potential (**Fig. 7c**), which is characteristic of hyperpolarization-activated cyclic nucleotide–gated (HCN) channel current (*I*_h_) expression^37^. Both voltage sag and voltage overshoot percentage were larger in L5 M1 neurons from PME mice (**Fig 7d**, see **Supplementary Table 4** for stats), although voltage overshoot did not quite reach the level of significance. Additional intrinsic properties of L5 M1 neurons in offspring can be viewed in **Supplementary Table 4**.

In addition to electrophysiological recordings, we measured the strength and organization of local synaptic inputs onto a single post-synaptic layer 5B M1 neuron by directing a UV laser beam (355 nm) focally uncage glutamate across a 16 x 16 stimulation grid centered around the recorded neuron (**Fig. 7b**). Compared to PSE (**Fig. 7g,h**), PME **(Fig. 7k,l**) modified the amplitude of local inputs onto layer 5 M1 neurons (see PME minus PSE amplitude difference images for females and males in **Fig 7i & m**, respectively). Row averages, which approximately correspond to subdivisions of cortex layers, were calculated by using the average amplitude of local inputs from rows 1:16, columns 6:13 of the stimulation grid. A significant interaction between exposure and row average was observed (rmANOVA: Exposure, F_(1,38)_=0.120, p=0.73; Row, F_(3.759,142.8)_=16.79, p<0.0001; Sex, F_(1,38)_=5.168, p=0.029; Row x Exposure, F_(15,570)_=2.411, p=0.0021; Row x Sex, F_(15,570)_=0.561, p=0.905; Exposure x Sex, F_(1,38)_=0.397, p=0.532; Row x Exposure x Sex, F_(15,570)_=1.126, p=0.329; **Fig. 7j,n**). In particular, increased amplitude of excitatory responses were observed following stimulation of local neurons in L2/3 (**Fig. 7i-j, m-n**). These electro-anatomical results suggest that the L2/3 → L5 local pathway is slightly enhanced in the motor cortex of juvenile PME mice. Representative morphological reconstruction of recorded neurons can be viewed in **Supplementary Fig. 14 a-b** (PSE and PME females, respectively) & **d-e** (PSE and PME males, respectively). Sholl analyses suggest that morphology of layer 5 M1 neurons in PME mice is unchanged from PSE littermates as neither intersections (rmANOVA: Exposure, F_(1,27)_=0.602, p=0.44; Radius, F_(16,432)_=148, p<0.0001; Sex, F_(1,27)_=7.34, p=0.012; Radius x Exposure, F_(16,432)_=0.632, p=0.86; Radius x Sex, F_(16,432)_=3.87, p<0.0001; Exposure x Sex, F_(1,27)_=0.0821, p=0.78; Radius x Exposure x Sex, F_(16,432)_=0.666, p=0.83; **Supplementary Fig. 14 c & f** *top*, for males and females, respectively) nor length (rmANOVA: Exposure, F_(1,27)_=1.09, p=0.30; Radius, F_(5.43,146.6)_= 119.6, p<0.0001; Sex, F_(1,27)_=6.56, p=0.016; Radius x Exposure, F_(16,432)_=0.659, p=0.83; Radius x Sex, F_(16,432)_=3.10, p<0.0001; Exposure x Sex, F_(1,27)_=0.305, p=0.59; Radius x Exposure x Sex, F_(16,432)_=0.586, p=0.89; **Supplementary Fig. 14 c & f** *bottom*, for males and females, respectively) were different between PME and PSE treatments.

**Figure 7.**
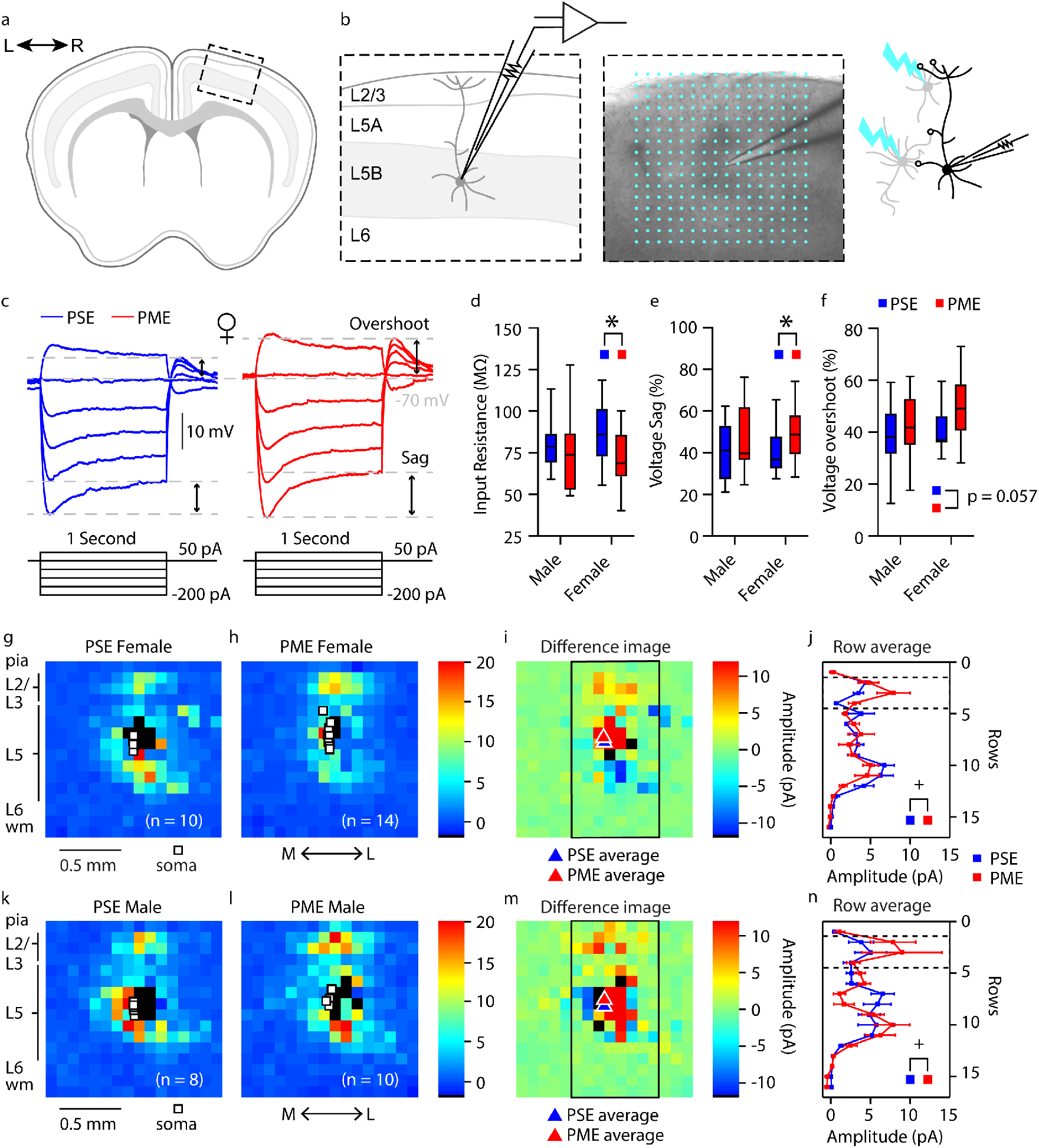
Intrinsic Properties and Local Circuit Mapping is Altered by PME. **a,b,** Schematic of electrophysiology recordings and circuit mapping. **a** Whole brains were extracted and coronally sectioned to acquire acute motor cortex slices for patch clamp electrophysiology. **b** Representation of a motor cortex neuron recording (left) and example (4X bright field video image) of a motor cortex neuron recording overlaid with the 16 × 16 stimulation grid (100μm spacing) for local circuit mapping (center). Schematic of laser scanning photo-stimulation of local neurons connected to recorded L5 pyramidal neuron in motor cortex (right). **c** Representative current-clamp traces of sub-threshold responses from L5 motor cortex neurons from PSE (blue traces) and PME (red traces) female mice. Step protocols and current values are displayed below traces. **d-f** PME decreased input resistance (**d**), increased voltage sag (**e**), and increased voltage overshoots (**f**) (ANOVA: Exposure, p=0.030, p=0.027, & p=0.057; n=13 PME mice (7M:6F), 41 cells (15M:26F) and n=11 PSE mice (6M:5F), 28 cells (15M:13F)). **g-h, k-l** Average local input maps from PSE (n=10 cells, 5 mice) and PME (n=14 cells, 4 mice) female (**g,h**) and PSE (n=8 cells, 5 mice) and PME (n=10 cells, 5 mice) male (**k,l**) mice. Each pixel represents the mean amplitude of the response evoked by UV photolysis of MNI-glutamate at that location. **i,m** PME mice demonstrate differences in local circuitry as identified in difference in PSE average map and PME average map for female (**i**) and male (**m**) mice. **j,n** This difference in local synaptic input was further supported by a significant Row X Exposure (rmANOVA: Interaction, p=0.0025) indicating PME altered local synaptic input on L5 M1 neurons in a region dependent manner (n=9 PME mice (5M:4F), 24 cells (10M:14F) and n=10 PSE mice (5M:5F), 18 cells (8M:10F). Row average data points indicate mean ± SEM where rows are the average input from rows 1:16, columns 6:13. Box plots indicate 25th to 75th percentiles with whisker characterizing the minimum and maximum value. * p<0.05 for main effect of exposure. + p<0.05 for row by exposure interaction.

## DISCUSSION

Given the rapid rise in infants born passively exposed to opioids during prenatal development, there is an urgent need to develop translationally relevant animal models of prenatal opioid exposure to begin to elucidate the impact of opioid exposure on development. The present study adds to the limited body of work examining preclinical models of prenatal opioid exposure. We provide the first longitudinal assessment of behavioral and neuronal development in offspring with PME combined with data on maternal and fetal opioid levels in the brain, placenta, and blood. Furthermore, the present study builds upon previous models by providing a more comprehensive, clinically relevant characterization of how methadone, a commonly used opioid during pregnancy, impacts development when offspring are opioid-exposed during the entire gestational period. Our findings reveal a disruption in physical, behavioral, and neuronal development in PME male and female offspring which persists beyond the immediate neonatal period and supports clinical studies that suggest babies born with opioid exposure are at higher risk for adverse developmental outcomes^9-11^.

Data on opioid levels in offspring are lacking in the preclinical literature. Bearing in mind differences in metabolism between rodents and humans, the methadone dose utilized here was likely in the clinical range^38,39^. Although dam plasma levels were lower that what is typically recommended for treating OUD in humans (~400 ng/mL)^40^, our dosing protocol produced opioid dependency in dams. Others also reported an accumulation of methadone in the rodent fetal brain^41^. We added to these findings by demonstrating methadone accumulated in the placenta and provided evidence that the placenta may have predictive power to infer the degree of prenatal brain opioid exposure during gestation. Although both methadone and EDDP accumulated in the placenta, EDDP did not concentrate in the fetal brain as methadone did. This observation may suggest the increase in methadone resulted from reduced metabolism in offspring. Methadone undergoes N-demethylation to form EDDP primarily via the cytochrome P450 enzyme CYP3A4 in humans^40^, but the expression of most cytochrome P450 enzymes in both mice and humans are dramatically lower during the prenatal period^42^. Nevertheless, we are confident that methadone was present in the fetal brain at pharmacologically-relevant levels in our model and likely contributed to the aberrant neurobehavioral development.

The persistent decrease in the size of offspring was striking, and other models of prenatal opioid exposure, including methadone, similarly demonstrated reduced body weight^13,17,18^. In humans, prenatal opioid exposure is similarly associated with low birth weight and being small for gestational age^43^, but it remains unclear what underlies this reduced size. Given the delayed acquisition of other sensorimotor milestones, it is possible PME also delayed acquisition of the suckling reflex which could contribute to reduced growth. An alternative hypothesis is that methadone disrupted the hypothalamic–pituitary–somatotropic axis. Central administration of exogenous opioids modulates plasma levels of growth hormone and insulin-like growth factor in adult humans and rodents^44,45^. Work done by Vathy & colleagues revealed that prenatal morphine exposure interacts with other hypothalamic-pituitary axes altering the homeostatic balance of stress-related hormones^46-48^, but the effect of prenatal opioid exposure on growth-related hormones has not been studied and may represent an important area of study for future work.

Separation-induced USVs emitted by mouse pups are frequently used as a measure of distress^27^. Production of these USVs is dependent on the endogenous opioid system. MOR agonists reduce USV rates, and mice lacking MORs do not vocalize when separated from the dam^49,50^. Others showed that PME reduces MOR binding affinity and alters downstream signaling^51^ suggesting the differences in USV development in our model resulted from changes in the endogenous opioid system. While the exact biological relevance of the increase in USV call rates and the alterations in call types and syntax on P7 is unclear at this time, these complex differences in USVs certainly reflected differential pup-dam communication between PME and PSE offspring.

PME offspring were hyperactive in the open field on P7 and P21 including a high number of jumps on P21. Combined with the increased USV call rate on P7, these activity findings are indicative of a heightened sensory arousal and hyperactive state in juvenile PME offspring. This hyperarousal phenotype may represent an enduring consequence of PME during the development of neurocircuitry controlling stress and/or sensorimotor activity. Our data support clinical findings that report higher risks of attention-deficit/hyperactivity disorder and increased hyperactivity and impulsivity in children with opioid exposure *in utero^25,26,52^.* In our model, PME offspring experienced delayed development of several sensorimotor milestones. Again, these findings mirror recent meta-analyses which support the negative impact of prenatal opioid exposure on psychomotor performance and motor function^9,10^. In regards to animal models, others also reported delayed development of certain sensorimotor milestones^12-14^, but these reports slightly differ in which specific milestones are delayed and which develop normally in offspring. For instance, offspring from our PME model did not exhibit differences in negative geotaxis, although in other models of prenatal buprenorphine and methadone exposure, negative geotaxis was disrupted^13,14^. Due to the differences in opioid used, duration and timing of exposure, and differences between rat and mouse development, identifying why specific milestones are delayed and why others are not remains challenging.

The neural mechanisms that could mediate the sensorimotor deficits and hyperactivity observed in our model are diverse. M1 has a central role in the preparation, execution, and adaptation of motor movements. Although the intrinsic properties of neurons can be affected by numerous ion channels and ion pumps, slice recording data indicated that altered intrinsic properties of M1 L5 pyramidal neurons from PME mice may be mediated by enhanced HCN channel-driven *I*_h_. In cortical pyramidal neurons, there is a gradient of HCN channel expression with the greatest being in the apical dendrites^53-55^. In mouse M1, HCN channels filter synaptic inputs onto L5 corticospinal neurons^37,56^. As a result, increased *I*_h_ in M1 L5 pyramidal neurons in PME mice suggests a narrowing of both spatial and temporal integration windows for both local and long-range synaptic inputs. Local circuit maps collected here showed that PME enhanced the L2/3 → L5 excitatory pathway in M1, which is the major local excitatory pathway for M1^56,57^. Therefore, it is possible that the narrowing of synaptic integration windows leads to compensatory mechanisms that enhanced the strength of synaptic inputs. These findings may suggest HCN channels represent a therapeutic target to prevent the potential neurobehavioral consequences of PME. In addition, methadone inhibits NMDA receptors^40^, which mediate experience-dependent synaptic pruning during maturation of the mouse cortex^58^. Therefore, PME-induced changes to dendritic pruning may also contribute to the altered local circuits of layer 5 M1 neurons, which could be further mediated by changes to HCN expression or function. While our morphological analysis revealed no effect of PME on L5 M1 neurons, measurements were highly variable indicating that neurons analyzed were a heterogeneous population of thick-tufted neurons.

As clinical evidence strongly indicates the use of opioid maintenance therapies for treating OUD improves pregnancy outcomes^4^, these findings which indicate PME impairs physical and neurobehavioral outcomes in offspring should not be interpreted as evidence against the use of opioid maintenance therapies in pregnancy. Instead, these data may suggest that early and preventative clinical intervention programs will be needed to support the healthy development of infants exposed to opioid *in utero.* Nonetheless, there are several limitations which must be considered when interpreting our findings. Prior reports indicated opioid treatment alters maternal behavior which could impact offspring development^14,59^. However, we and others^60-62^, did not observe differences in maternal care suggesting the physical, behavioral, and neuronal differences found in offspring are a result of passive methadone exposure during gestational development. Nonetheless, we cannot be confident that subtle differences in maternal behavior interacted with methadone exposure to alter offspring outcomes. Next, we modeled PME via noncontingent treatment using twice daily injections. Although free-access or self-administration would be more translational, noncontingent exposure reduces variability in methadone exposure between pregnant females. The interaction of PME with the stress of injections and repeated handling of dams could produce the differential offspring outcomes. Given that medically supervised opioid withdrawal is not recommended for OUD, our choice of prenatal opioid exposure model was based on epidemiological trends which supports the high usage of methadone by pregnant women^19^. However, there exists numerous variations in human opioid use during pregnancy which our model does not reflect (e.g. women may not initiate opioid maintenance therapy until partway through pregnancy or they may experience a relapse and use other recreational opioids at variable intervals during pregnancy). Further work by our group and others will be required to examine these variants of prenatal opioid exposure. The differences between rodent and human gestational development are an inherent limitation of our model that cannot be overcome. The first week of rodent postnatal life is roughly equivalent to the third trimester^12^. In an attempt to maintain a sufficient level of methadone exposure to the newborn pups during P0-P7, the dams were maintained at their highest gestational dose after giving birth and up to P7; however, it is unlikely this strategy completely replicates *in utero* third trimester opioid exposure in humans. Lastly, our findings in offspring were collected in the preweaning and early adolescent periods. Future studies will examine if alterations in physical, behavioral, and neuronal outcomes persist into adulthood or if PME offspring are able to eventually overcome these differences.

## METHODS

### Animals and Model Generation

Animal care and research were conducted in accordance with guidelines established by the National Institutes of Health and protocols were approved by the Indiana University School of Medicine Institutional Animal Care and Use Committee. Eight-week-old female C57BL/6J mice were acquired from Jackson Laboratories (Bar Harbor, Maine), single housed, and randomly assigned to either saline (10 mL/kg) or oxycodone treatments. Oxycodone dependence was induced by a dose-ramping procedure with a dose of 10 mg/kg administered on pregestational day (PG) 14, 20 mg/kg on PG13, and then maintained on 30 mg/kg for PG12-6. All saline or oxycodone doses were administered subcutaneously twice daily at least seven hours apart. On PG5, oxycodone-treated mice were transitioned to 10 mg/kg methadone (s.c. b.i.d.) while saline-treated animals continued to receive saline injections. Oxycodone and methadone were obtained from the National Institute on Drug Abuse Drug Supply Program. Five days following initiation of methadone treatment, an 8-week-old C57BL/6J male mouse (also acquired from Jackson Laboratories) was placed into the cage of each female for four days. Mucous plugs were assessed each morning to approximate gestational day (G) 0. Food consumption was examined weekly in a subset of these female mice. Female mice were weighed every Monday, Wednesday, and Friday throughout the study with the exception of the mating period.

Cages were examined for the presence of pups at the time of each morning and afternoon injection, and the day of birth was designated postnatal day (P) 0. Data on litter size, sexes, and neonatal deaths were collected. The presence of unconsumed placentas at 24 hours following birth was also examined. Only litters between three and eight pups were used in subsequent studies of offspring. In an attempt to maintain a sufficient level of methadone exposure to the newborn pups during P0-P7, the dams were maintained at their highest gestational dose after giving birth and up to P7. After P7, the dose of methadone administered to dams was adjusted to their body weight.

### Methadone and Metabolite Concentrations

All tissue and blood samples were collected 2.5 hours following morning administration of methadone. On G18, three dams were anesthetized with ketamine/xylazine and fetuses were rapidly removed from the uterine cavity to examine brain and placental drug levels. On P1 and P7, three dams and their litters were sacrificed for the collection of trunk blood and brains. Blood samples were centrifuged for five minutes at 4,500g to collect 20 uL of plasma. Plasma, placenta, and whole brains were frozen in isopentane on dry ice and stored in −80°C until processing. The samples were examined for the presence of methadone and EDDP via high performance liquid chromatography tandem mass spectroscopy (HPLC–MS/MS; Sciex 5500 QTRAP, Applied Biosystems, Foster City, MA). The analytical method for plasma samples has previously been described^63^. Tissue samples and standards were prepared by adding diphenhydramine (internal standard, 0.1 ng/μL in methanol) and 0.1 M phosphate buffer (200 μL, pH 7.4) to tissue samples. Samples were then extracted using a liquid-liquid procedure with 3 mL ethyl acetate. The supernatant was transferred to a clean tube and evaporated to dryness, then reconstituted in acetonitrile with 0.1% formic acid (pH 6.5). Ten microliters of sample were then injected and the remaining protocol follows the previously published procedure^63^. The limit of quantification for methadone and EDDP detection was 0.1 ng/mL and 0.05 ng/mL in the plasma, respectively, and 0.08 ng/sample and 0.04 ng/sample of placenta and brain, respectively for methadone and EDDP.

### Maternal Characteristics

Dams were given a new compressed paper nestlet, and twelve hours later, the ability of dams to build a new nest on P3 was assessed using a criterion described elsewhere^64^. A pup retrieval task was completed by first removing one pup and the dam and then placing the pup at one end of the cage (~30 cm away). The dam was then released at the other end of the cage and the time required to retrieve the missing pup was recorded. This was repeated with two other pups and an average pup retrieval latency score was calculated.

To demonstrate that the oxycodone dosing strategy induced dependency, a subset of mice underwent naloxone-precipitated withdrawal prior to methadone transition. Sixty to ninety minutes following the final 30 mg/kg oxycodone dose on PG6, all mice were administered naloxone (5 mg/kg, i.p.) and somatic signs of withdrawal were assessed for ten minutes using a previously described protocol^65^. These mice which underwent precipitated withdrawal were not used in any subsequent studies. Similarly, following the weaning of offspring, a subset of dams underwent the naloxone-precipitated withdrawal procedure to demonstrate that the transition to the 10 mg/kg methadone dosing regimen maintained opioid dependency.

### Offspring Physical Development

Beginning at P0 and throughout the pre-weaning period, offspring were marked for identification on the paws with a red or black marker. The progression of weight gain and body length on P0, P3, P7, P10, P14, P18, and P21 was examined. A subset of the offspring was also weighed at P35 and P49. From P3-P18, offspring were examined for the first day both eyes were open, the first day both ear pinnae completely unfolded and the ear canal was patent, and the first day the lower incisors emerged.

The right femur of P7 and P35 offspring was scanned on a SkyScan 1172 (Bruker, Billerica, MA, USA) with a 0.5 aluminum filter and a 6 μm voxel size. For P7 samples, mineralized femur length was obtained from a coronal view of the scan. Whole bone volume was obtained in the transaxial plane incorporating the entire mineralized bone. For P35 samples, a 0.5 mm region of interest at the distal femur was selected proximal to the growth plate. Distal femur metaphysis bone volume was obtained in the transaxial plane from this whole region including both trabecular and cortical bone. Trabecular bone was isolated in the same region for trabecular bone volume. Cortical bone in P35 samples was measured from five slices 2 mm proximal the end of the metaphysis region of interest. Cortical bone area and cortical thickness were measured at this site.

### Offspring Behavior

#### Open Field

A modified open field (12.5 W x 15.8 L x 13.7 H cm) with a video recording device (Brio Ultra HD Pro Webcam, Logitech, Lausanne, Switzerland) and an ultrasound microphone (Pettersson Elektronik AB, Uppsala, Sweden) was used to assess locomotion and USVs development for five minutes on P1, P7, P14, and P21 in offspring. These sessions began at 8:00 A.M following a one-hour acclimation period. The arena was placed on a far infrared warming pad (Kent Scientific, Torrington, CT) that maintained the surface temperature between 35-37°C. The entire apparatus was located within an environmental control chamber (Omnitech Electronics Inc., Columbus, OH) that was layered with sound proofing material. USV audio files were processed using the open-source, deep learning-based system DeepSqueak^28^. Video tracking analysis was accomplished with EthoVision XT software (Noldus, Leesburg, VA). P1 recorded videos were examined for the number of twitches/jerks, a behavior consistent with NOWS in humans and rodents^23,66^. As temperature instability is also a symptom of NOWS^23^, body temperature was assessed on P1. Without a heating source, body temperature was monitored in offspring at baseline and after two minutes of isolation. Due to the small size of pups preventing collection of rectal temperatures, a FLIR E8 infrared camera (FLIR Systems, Wilsonville, OR) was used to capture surface body temperature. Post-hoc analysis of duration of time spent near the walls of the arena was assessed using Noldus software. Noldus Automatic Behavior Recognition was used to track the frequency of jumping, unsupported and supported rearing, and grooming behavior.

#### Sensorimotor Milestones

Beginning on P3 and through P14, offspring were tested daily to examine acquisition of: surface righting to prone position within five seconds of release, expression of negative geotaxis within 15 seconds of release, display of cliff aversion within ten seconds of release on a ledge, ability to grasp a suspended rod with both forelimbs for at least two seconds, and extinguishing of pivoting behavior demonstrated by walking outside of a 12 cm diameter circle within 30. These assessments are well-established paradigms used to characterize the normal development of newborn mice^67^. These sessions began at 1:00 P.M. following a one-hour acclimation period. Acquisition of these sensorimotor developmental milestones was considered complete by a successful demonstration of the behavior within the allotted time for three consecutive days in a row.

#### Acoustic Startle and Prepulse Inhibition

Acoustic startle responses and PPI were assessed in P28-P29 mice in isolated chambers (SR-LAB™ Startle Response System, SD Instruments, San Diego, CA). Following five 115 Db startle pulses for acclimation, mice were exposed to 8 blocks consisting of either a single pulse at the background level, 65, 75, 85, 95, 105, or 115 Db to assess acoustic startle response or a prepulse of either 65, 75, or 85 Db followed by a 115 Db startle pulse to assess PPI in random order. The intertrial interval was randomly assigned with intervals between 10-40 seconds. An accelerometer detected changes in force due to jumping/flinching and the output was an excitation voltage change in millivolts (mV).

### Magnetic Resonance Imaging

Volumetric MRI studies were performed using a 9.4 T Bruker system (Bruker BioSpin MRI GmbH, Germany) equipped with a BGA-12S gradient. A Bruker 2-channel surface mouse head coil was used for high resolution brain imaging. Animals were anesthetized and maintained with 1.5-2% isoflurane during MRI sessions. T2-weighted images with a three-dimensional Rapid Acquisition with Relaxation Enhancement (RARE) sequence were acquired for volumetric measurements (TR/TE=1500/11 ms, Rare Factor=12, FOV=20 x 20 x 7 mm with an isotropic voxel of 120 μm^3^, number of averages=1, scan time=11.8 min).

A mouse brain atlas for 4-week-old mice was created by modifying PMOD (PMOD Technologies LLC, Switzerland) Ma-Benveniste-Mirrione mouse brain atlas. Specifically, the T2-weighted MRI images of a 4-week-old control mouse were normalized to the PMOD built-in mouse brain template using a Deform function in the ‘Fuse It’ module. VOIs were then manually adjusted to fit the 4-week-old mouse brain and to create a 4-week mouse atlas for this study. To obtain the brain volume measurements, T2-weighted anatomical images from individual mouse in the native space were normalized to the 4-week mouse template and an inverse transformation matrix was saved in Fuse It module. The native space image data were loaded again in the View module then 4-week-mouse atlas with VOIs was loaded and inverse transformed to obtain the volume measurements. (See **Supplementary Fig. 8** for segmentation).

### Immunostaining

On P22-24, offspring were anesthetized with isoflurane and perfused with 4% paraformaldehyde prepared in PBS for ten minutes at a pump rate of ~2 mL/minute. Fixed brains were sectioned into 100 μm sections in the coronal plane using a Leica VT-1000 vibrating microtome (Leica Microsystems) and stored in PBS until later analysis. Sections were permeabilized with 0.3% Triton X100, then incubated with a blocking solution (3% normal goat serum prepared in PBS with 0.3% Triton X-100) and then incubated overnight with primary antibody (see **Supplementary Table 3**) prepared in blocking solution. An appropriate secondary antibody conjugated with an Alexa series fluorophore (see **Supplementary Table 3**) was used to detect the primary antibody. Draq5 (1:10,000 dilution, Cell Signaling) was included in the secondary antibody solution to stain nuclei. Z-stack confocal images were acquired from both hemispheres with a Leica confocal microscope with a 10X/NA0.75 objective. The Z-stacks were taken at 0.5 μm intervals, 5 μm-total thickness was imaged. Projection images of 5 μm-thickness were used for image quantification by using NIH ImageJ software. The regions of interest were defined as bins 1-10 for the ACC and S1 and defined at layer II-IV and layer 5 in M1.

### Electrophysiology, Mapping, Morphology, Imaging

#### Electrophysiology and mapping

At P21-26, acute brain slices were prepared as described previously^37,68^. Pipettes for recordings were fabricated from borosilicate capillaries with filaments (G150-F, Warner) using a horizontal puller (P-97, Sutter). For acquiring the electrophysiological profiles of neurons, patch pipettes contained potassium-based intracellular solution (in mM: 128 K-gluconate, 10 HEPES, 1 EGTA, 4 MgCl2, 4 ATP, and 0.4 GTP, 10 phosphocreatine, 3 ascorbate; pH 7.2). 3-4 mg biocytin (Sigma-Aldrich) was included in the intrapipette solution for morphological reconstruction. MNI-caged glutamate (Tocris Bioscience, 295325-62-1) [0.2 mM] was combined with recirculating solution for all recordings. The bath solution for recordings contained elevated concentrations of divalent cations [4 mM Ca^2+^ and 4 mM Mg^2+^] and NMDA receptor antagonist 3-(2-Carboxypiperazin-4-yl)propyl-1-phosphonic acid (CPP) [5 mM] (Tocris biosciences, 126453-07-4). Slices were used 1.5–4.5 hours after preparation. Recordings were performed at room temperature. Pipette series resistance was between 2-5 MΩ. Pipette capacitance was compensated; series resistance was monitored but not compensated and required to be <25 MΩ for inclusion in the data set. Current-clamp recordings were bridge-balanced. Current was injected as needed to maintain the membrane potential near −70 mV during select stimulus protocols. Recordings were filtered at 4 kHz and sampled at 10 kHz using a Multiclamp 700B amplifier (Molecular Devices). Membrane potential values were not corrected for a calculated liquid junction potential of 11 mV. Ephys software (http://www.ephus.org) was used for hardware control and data collection^69^. Methods for determining input resistance, voltage threshold for action potential (AP) firing, and voltage sag have been reported previously^36^. Glutamate uncaging and laser scanning photo-stimulation (glu-LSPS) were performed as described previously^70^. Once a patch recording of a labeled neuron was established, an image of the slice (4X objective) was acquired before mapping for precise registration of the mapping grid. The mapping grid (16×16; 100 μm spacing) was rotated with the top row of the grid flush with the exterior primary motor area layer one. The grid locations were sampled (every 0.4 s) with a UV stimulus 1.0 ms in duration and 20 mW at the specimen plane. Photo-stimulation sites resulting in activation of glutamate receptors on the dendrites of the recorded neuron were readily detected based on characteristically short onset latencies (7 ms) of responses^56^ and included in the map analyses as synaptic responses resulting from uncaged glutamate activation of presynaptic neurons within the local circuit. Responses overlapped by direct activation of the recorded neuron’s dendrites were excluded and rendered as black pixels in input color maps. These maps thus represent “images” of the local sources of monosynaptic input arising from small clusters of ~100 neurons at each stimulus location. Excitatory (glutamatergic) responses were recorded at a command voltage of −70 mV. Excitatory input maps were constructed on the basis of the mean inward current over a 0–50 ms post-stimulus time window.

#### Morphology

Following electrophysiology recordings, the pipette was removed slowly at an angle in voltage clamp mode while monitoring the capacitive transients in order to reestablish a seal. Following re-sealing of the final cell in a slice, the slice was removed from the recording chamber and left in oxygenated ACSF solution for up to 3 hours, to ensure transport of biocytin to distal processes. The slice was then placed in 4% paraformaldehyde solution in 0.1 M Phosphate Buffer (4% PFA/PB) for 24 hours. Slices were washed (3x for 5 minutes at room temperature while shaking) with 0.1 M PBS and 0.1% Triton-X-100 (Acros Organics, AC21568-2500), then nutated in blocking solution containing normal goat serum (Millipore Sigma, 566380) for one hour. Slices were washed again, then nutated in blocking solution with secondary antibody Streptavidin Alexafluor conjugate 488 (1:1000; Fisher Scientific, S32354) at room temperature for one hour. Slices were washed and mounted to glass slides. Images of recovered biocytin-filled neurons were taken using a Nikon A1R confocal microscope with x40 oil immersion objective. Z-stack images were taken at a pitch of 1.5 μm and stitched together with 15% overlap. Neurolucida and Neurolucida 360 (MBF Bioscience) was used for reconstruction and Sholl analysis.

#### Data Exclusion Criteria

10/125 neurons were excluded from analysis due to pipette series resistance >25 MΩ and instability over time, 6 of which were PME treated neurons from male mice. After morphological reconstruction and image acquisition, additional inclusion criteria were established: 1) imaged M1 neurons with no pia-directed dendrite were excluded from all analyses (6/115), 2) imaged motor cortex neurons with no apical tuft at the end of a pia-directed dendrite were excluded from excitatory input map analysis on the assumption that vibratome slicing separated apical tuft from cell body (8/115) and 3) M1 neurons which did not reseal and were unable to be imaged (5/115), or neurons which did not have ‘maps’ collected (11/115), were excluded from excitatory input map analysis. 20/115 neurons were not processed for morphological reconstruction but were included in electrophysiology intrinsic data analysis. Thin-tufted neurons (23) were excluded from analysis on the basis of <20% voltage sag. 42 thick-tufted neurons were utilized for map analysis and of these, 32 were reconstructed for morphological analysis.

### Statistics

Experimenters were blinded to treatment/exposure group for data collection of all studies. Statistical analyses were conducted using GraphPad Prism 8 (San Diego, CA). Data are graphically presented as the mean ± SEM or as box plots extending from the 25th to 75th percentiles with whisker characterizing minimum and maximum values. The level of significance was set at p<0.05. All experiments were performed using both male and female offspring. To minimize potential litter effects in all completed studies, offspring from 4 or more litters per exposure were utilized as previously done^16^. For offspring weight and length, offspring from 8-9 litters per exposure were examined. For all behavioral studies, offspring were taken from 4-5 different litters per exposure. For neuroimaging and immunohistochemistry studies, offspring from 4 litters per exposure were utilized. For electrophysiology, mapping, morphology, and imaging, neurons were recorded and imaged from offspring of 9 different litters per exposure. With the exception of MRI and immunostaining, all studies were sufficiently powered to detect sex differences. However, for clarity of focusing on the effect of prenatal exposure, when no main effects of sex or interactions with sex were found in offspring, the data was collapsed on sex and re-analyzed. Chi-square tests for categorical data was used for litter characteristics. A linear regression was performed to assess the relationship between placental methadone levels and offspring brain levels on G18, and offspring plasma and brain levels on P1 and P7. Two-tailed unpaired t-tests were used to analyze normally distributed data and Mann-Whitney U tests or Wilcoxon rank sum test (for electrophysiology data only) were used to analyze non-normally distributed data. For data with multiple groups and/or repeated measures, ANOVAs with Sidak’s post hoc tests were used.

## Funding

This work was supported by grants awarded by the National Institutes of Health R01AA027214 (BKA), T32AA07462 (KCR, DLH), F30AA028687 (GGG), Indiana University (BKA, BKY, HCL), Indiana University Health (BKA), the Stark Neurosciences Research Institute (BKA, DLH, GGG), IU Simon Cancer Center Support Grant P30CA082709 (ARM).

## Acknowledgments

We thank the National Institute on Drug Abuse Drug Supply Program for generously providing the methadone and oxycodone utilized in the experiments of this manuscript. Mass spectrometry work was provided by the Clinical Pharmacology Analytical Core at Indiana University School of Medicine; a core facility supported by the IU Simon Cancer Center Support Grant P30 CA082709. We thank Dr. David McKinzie, the director of the Behavioral Phenotyping Core at Indiana University School of Medicine, for valuable comments on study design.

## Author Contributions

GGG, BM, HJH, CEM, DLH, CWM, JK, BKY, MRA, YCW, PLS, BKA designed experiments. GGG, BM, HJH, CEM, DLH, KCR, YG, HH, SNK, ARM, CWM, and EAN performed experiments and contributed to data analyses. All authors discussed the results and contributed to all stages of manuscript preparation and editing.

## Competing Interests

The authors have no competing interests to report.

## Notes

### Competing Interest Statement

The authors have declared no competing interest.

### Summary of Updates

This revision contains updates to clarify some introductory information on the model, further additions of limitations to the work, and additional details of methods.

## References

1 Haight, S.C., Ko, J. Y., Tong, V. T., Bohm, M.K. & Callaghan, W. M. Opioid Use Disorder Documented at Delivery Hospitalization - United States, 1999-2014. MMWR. Morbidity and mortality weekly report 67, 845–849, doi:10.15585/mmwr.mm6731a1 (2018).

2 Ko, J. Y. et al. Incidence of Neonatal Abstinence Syndrome - 28 States, 1999-2013. MMWR. Morbidity and mortality weekly report 65, 799–802, doi:10.15585/mmwr.mm6531a2 (2016).

3 Patrick, S. W., Davis, M. M., Lehmann, C. U. & Cooper, W. O. Increasing incidence and geographic distribution of neonatal abstinence syndrome: United States 2009 to 2012. Journal of perinatology: official journal of the California Perinatal Association 35, 650–655, doi:10.1038/jp.2015.36 (2015).

4 Practice, A. C. o. O. Opioid Use and Opioid Use Disorder in Pregnancy: ACOG Committee Opinion No. 711. Obstetrics & Gynecology 130, 81–94 (2017).

5 Towers, C. V. et al. Neonatal Head Circumference in Newborns With Neonatal Abstinence Syndrome. Pediatrics 143, doi:10.1542/peds.2018-0541 (2019).

6 Sirnes, E. et al. Brain morphology in school-aged children with prenatal opioid exposure: A structural MRI study. Early human development 106-107, 33–39, doi:10.1016/j.earlhumdev.2017.01.009 (2017).

7 Hartwell, M. L., Croff, J. M., Morris, A. S., Breslin, F. J. & Dunn, K. Association of Prenatal Opioid Exposure With Precentral Gyrus Volume in Children. JAMA Pediatrics, doi:10.1001/jamapediatrics.2020.0937 (2020).

8 Monnelly, V. J. et al. Prenatal methadone exposure is associated with altered neonatal brain development. NeuroImage. Clinical 18, 9–14, doi:10.1016/j.nicl.2017.12.033 (2018).

9 Yeoh, S. L. et al. Cognitive and Motor Outcomes of Children With Prenatal Opioid Exposure: A Systematic Review and Meta-analysis. JAMA network open 2, e197025, doi:10.1001/jamanetworkopen.2019.7025 (2019).

10 Lee, S. J., Bora, S., Austin, N., Westerman, A. & Henderson, J. M. T. NEURODEVELOPMENTAL OUTCOMES OF CHILDREN BORN TO OPIOID-DEPENDENT MOTHERS: A SYSTEMATIC REVIEW AND META-ANALYSIS. Academic pediatrics, doi:10.1016/j.acap.2019.11.005 (2019).

11 Larson, J. J. et al. Cognitive and Behavioral Impact on Children Exposed to Opioids During Pregnancy. Pediatrics, doi:10.1542/peds.2019-0514 (2019).

12 Robinson, S. A., Jones, A. D., Brynildsen, J. K., Ehrlich, M. E. & Blendy, J. A. Neurobehavioral effects of neonatal opioid exposure in mice: Influence of the OPRM1 SNP. Addiction Biology 0, e12806, doi:10.1111/adb.12806 (2019).

13 Kunko, P. M. et al. Perinatal methadone exposure produces physical dependence and altered behavioral development in the rat. The Journal of pharmacology and experimental therapeutics 277, 1344–1351 (1996).

14 Wallin, C. M., Bowen, S. E., Roberge, C. L., Richardson, L. M. & Brummelte, S. Gestational buprenorphine exposure: Effects on pregnancy, development, neonatal opioid withdrawal syndrome, and behavior in a translational rodent model. Drug and Alcohol Dependence 205, 107625, doi:https://doi.org/10.1016/j.drugalcdep.2019.107625 (2019).

15 Lu, R., Liu, X., Long, H. & Ma, L. Effects of prenatal cocaine and heroin exposure on neuronal dendrite morphogenesis and spatial recognition memory in mice. Neuroscience letters 522, 128–133, doi:10.1016/j.neulet.2012.06.023 (2012).

16 Ricalde, A. A. & Hammer, R. P., Jr. Perinatal opiate treatment delays growth of cortical dendrites. Neuroscience letters 115, 137–143, doi:10.1016/0304-3940(90)90444-e (1990).

17 Jantzie, L. L. et al. Prenatal opioid exposure: The next neonatal neuroinflammatory disease. Brain, Behavior, and Immunity, doi:https://doi.org/10.1016/j.bbi.2019.11.007 (2019).

18 Byrnes, E. M. & Vassoler, F. M. Modeling prenatal opioid exposure in animals: Current findings and future directions. Frontiers in neuroendocrinology 51, 1–13, doi:10.1016/j.yfrne.2017.09.001 (2018).

19 Duffy, C. R. et al. Trends and Outcomes Associated With Using Long-Acting Opioids During Delivery Hospitalizations. Obstet Gynecol 132, 937–947, doi:10.1097/AOG.0000000000002861 (2018).

20 Yazdy, M. M., Desai, R. J. & Brogly, S. B. Prescription Opioids in Pregnancy and Birth Outcomes: A Review of the Literature. J Pediatr Genet 4, 56–70, doi:10.1055/s-0035-1556740 (2015).

21 Mattia, C., Di Bussolo, E. & Coluzzi, F. Non-analgesic effects of opioids: the interaction of opioids with bone and joints. Current pharmaceutical design 18, 6005–6009, doi:10.2174/138161212803582487 (2012).

22 Mullens, C. L., McCulloch, I. L., Hardy, K. M., Mathews, R. E. & Mason, A. C. Associations between Orofacial Clefting and Neonatal Abstinence Syndrome. Plast Reconstr Surg Glob Open 7, e2095–e2095, doi:10.1097/GOX.0000000000002095 (2019).

23 Kocherlakota, P. Neonatal Abstinence Syndrome. 134, e547–e561, doi:10.1542/peds.2013-3524 %J Pediatrics (2014).

24 Andersen, J. M., Høiseth, G. & Nygaard, E. Prenatal exposure to methadone or buprenorphine and long-term outcomes: A meta-analysis.Early human development 143, 104997, doi:https://doi.org/10.1016/j.earlhumdev.2020.104997 (2020).

25 Azuine, R. E. et al. Prenatal Risk Factors and Perinatal and Postnatal Outcomes Associated With Maternal Opioid Exposure in an Urban, Low-Income, Multiethnic US Population. JAMA network open 2, e196405, doi:10.1001/jamanetworkopen.2019.6405 (2019).

26 Sundelin Wahlsten, V. & Sarman, I. Neurobehavioural development of preschool-age children born to addicted mothers given opiate maintenance treatment with buprenorphine during pregnancy. Acta paediatrica (Oslo, Norway: 1992) 102, 544–549, doi:10.1111/apa.12210 (2013).

27 Ehret, G. Infant rodent ultrasounds-a gate to the understanding of sound communication. Behavior genetics 35, 19–29 (2005).

28 Coffey, K. R., Marx, R. G. & Neumaier, J. F. DeepSqueak: a deep learning-based system for detection and analysis of ultrasonic vocalizations. Neuropsychopharmacology 44, 859–868, doi:10.1038/s41386-018-0303-6 (2019).

29 Frau, R. et al. Prenatal THC exposure produces a hyperdopaminergic phenotype rescued by pregnenolone. Nature Neuroscience 22, 1975–1985, doi:10.1038/s41593-019-0512-2 (2019).

30 Paus, T. Primate anterior cingulate cortex: Where motor control, drive and cognition interface. Nature Reviews Neuroscience 2, 417–424, doi:10.1038/35077500 (2001).

31 Hatsopoulos, N. G. & Suminski, A. J. Sensing with the motor cortex. Neuron 72, 477–487, doi:10.1016/j.neuron.2011.10.020 (2011).

32 Umeda, T., Isa, T. & Nishimura, Y. The somatosensory cortex receives information about motor output. Science Advances 5, eaaw5388, doi:10.1126/sciadv.aaw5388 (2019).

33 Hattox, A. M. & Nelson, S. B. Layer V neurons in mouse cortex projecting to different targets have distinct physiological properties. J Neurophysiol 98, 3330–3340 (2007).

34 Morishima, M., Morita, K., Kubota, Y. & Kawaguchi, Y. Highly differentiated projection-specific cortical subnetworks. The Journal of neuroscience: the official journal of the Society for Neuroscience 31, 10380–10391, doi:10.1523/JNEUROSCI.0772-11.2011 (2011).

35 Oswald, M. J., Tantirigama, M. L., Sonntag, I., Hughes, S. M. & Empson, R. M. Diversity of layer 5 projection neurons in the mouse motor cortex. Frontiers in cellular neuroscience 7, 174, doi:10.3389/fncel.2013.00174 (2013).

36 Suter, B. A., Migliore, M. & Shepherd, G. M. Intrinsic electrophysiology of mouse corticospinal neurons: a class-specific triad of spike-related properties. Cerebral cortex (New York, N.Y.: 1991) 23, 1965–1977, doi:10.1093/cercor/bhs184 (2013).

37 Sheets, P. L. et al. Corticospinal-specific HCN expression in mouse motor cortex: I(h)-dependent synaptic integration as a candidate microcircuit mechanism involved in motor control. J Neurophysiol 106, 2216–2231, doi:10.1152/jn.00232.2011 (2011).

38 Devidze, N. et al. Steady-state methadone effect on generalized arousal in male and female mice. Behav Neurosci 122, 1248–1256, doi:10.1037/a0013276 (2008).

39 Dryden, C., Young, D., Hepburn, M. & Mactier, H. Maternal methadone use in pregnancy: factors associated with the development of neonatal abstinence syndrome and implications for healthcare resources. BJOG: an international journal of obstetrics and gynaecology 116, 665–671, doi:10.1111/j.1471-0528.2008.02073.x (2009).

40 Eap, C. B., Buclin, T. & Baumann, P. Interindividual variability of the clinical pharmacokinetics of methadone: implications for the treatment of opioid dependence. Clinical pharmacokinetics 41, 1153–1193, doi:10.2165/00003088-200241140-00003 (2002).

41 Kongstorp, M., Bogen, I. L., Stiris, T. & Andersen, J. M. High Accumulation of Methadone Compared with Buprenorphine in Fetal Rat Brain after Maternal Exposure. The Journal of pharmacology and experimental therapeutics 371, 130–137, doi:10.1124/jpet.119.259531 (2019).

42 Hart, S. N., Cui, Y., Klaassen, C. D. & Zhong, X.-b. Three patterns of cytochrome P450 gene expression during liver maturation in mice. Drug Metab Dispos 37, 116–121, doi:10.1124/dmd.108.023812 (2009).

43 Nørgaard, M., Nielsson, M. S. & Heide-Jørgensen, U. Birth and Neonatal Outcomes Following Opioid Use in Pregnancy: A Danish Population-Based Study. Subst Abuse 9, 5–11, doi:10.4137/SART.S23547 (2015).

44 Hashiguchi, Y., Molina, P. E., Fan, J., Lang, C. H. & Abumrad, N. N. Central opiate modulation of growth hormone and insulin-like growth factor-I. Brain Research Bulletin 40, 99–104, doi:https://doi.org/10.1016/0361-9230(96)00045-7 (1996).

45 Abs, R. et al. Endocrine consequences of long-term intrathecal administration of opioids. The Journal of clinical endocrinology and metabolism 85, 2215–2222, doi:10.1210/jcem.85.6.6615 (2000).

46 Vathy, I. Prenatal opiate exposure: long-term CNS consequences in the stress system of the offspring. Psychoneuroendocrinology 27, 273–283 (2002).

47 Slamberová, R., Rimanóczy, A., Riley, M. A. & Vathy, I. Hypothalamo-pituitary-adrenal axis-regulated stress response and negative feedback sensitivity is altered by prenatal morphine exposure in adult female rats. Neuroendocrinology 80, 192–200, doi:10.1159/000082359 (2004).

48 Rimanóczy, A., Slamberová, R., Riley, M. A. & Vathy, I. Adrenocorticotropin stress response but not glucocorticoid-negative feedback is altered by prenatal morphine exposure in adult male rats. Neuroendocrinology 78, 312–320, doi:10.1159/000074884 (2003).

49 Carden, S. E., Barr, G. A. & Hofer, M. A. Differential effects of specific opioid receptor agonists on rat pup isolation calls. Brain research. Developmental brain research 62, 17–22, doi:10.1016/0165-3806(91)90185-l (1991).

50 Moles, A., Kieffer, B. L. & D’Amato, F. R. Deficit in attachment behavior in mice lacking the mu-opioid receptor gene. Science (New York, N.Y.) 304, 1983–1986, doi:10.1126/science.1095943 (2004).

51 Kongstorp, M. et al. Prenatal exposure to methadone or buprenorphine alters μ-opioid receptor binding and downstream signaling in the rat brain. International Journal of Developmental Neuroscience n/a, doi:10.1002/jdn.10043.

52 Ornoy, A., Segal, J., Bar-Hamburger, R. & Greenbaum, C. Developmental outcome of school-age children born to mothers with heroin dependency: importance of environmental factors. Developmental medicine and child neurology 43, 668–675, doi:10.1017/s0012162201001219 (2001).

53 Berger, T., Larkum, M. E. & Luscher, H. R. High I(h) channel density in the distal apical dendrite of layer V pyramidal cells increases bidirectional attenuation of EPSPs. J Neurophysiol 85, 855–868 (2001).

54 Magee, J. C. Dendritic hyperpolarization-activated currents modify the integrative properties of hippocampal CA1 pyramidal neurons. The Journal of neuroscience: the official journal of the Society for Neuroscience 18, 7613–7624 (1998).

55 Stuart, G. & Spruston, N. Determinants of voltage attenuation in neocortical pyramidal neuron dendrites. The Journal of neuroscience: the official journal of the Society for Neuroscience 18, 3501–3510. (1998).

56 Anderson, C. T., Sheets, P. L., Kiritani, T. & Shepherd, G. M. Sublayer-specific microcircuits of corticospinal and corticostriatal neurons in motor cortex. Nat Neurosci 13, 739–744, doi:10.1038/nn.2538 (2010).

57 Weiler, N., Wood, L., Yu, J., Solla, S. A. & Shepherd, G. M. Top-down laminar organization of the excitatory network in motor cortex. Nat Neurosci 11, 360–366 (2008).

58 Yu, X. et al. Accelerated experience-dependent pruning of cortical synapses in ephrin-A2 knockout mice. Neuron 80, 64–71, doi:10.1016/j.neuron.2013.07.014 (2013).

59 Slamberova, R., Szilagyi, B. & Vathy, I. Repeated morphine administration during pregnancy attenuates maternal behavior. Psychoneuroendocrinology 26, 565–576, doi:10.1016/s0306-4530(01)00012-9 (2001).

60 Alipio, J. B. et al. Enduring consequences of perinatal fentanyl exposure in mice. Addict Biol, e12895, doi:10.1111/adb.12895 (2020).

61 Tan, J. W. et al. Impaired contextual fear extinction and hippocampal synaptic plasticity in adult rats induced by prenatal morphine exposure. Addict Biol 20, 652–662, doi:10.1111/adb.12158 (2015).

62 Kongstorp, M., Bogen, I. L., Stiris, T. & Andersen, J. M. Prenatal exposure to methadone or buprenorphine impairs cognitive performance in young adult rats. Drug and Alcohol Dependence 212, 108008, doi:https://doi.org/10.1016/j.drugalcdep.2020.108008 (2020).

63 Metzger, I. F. et al. Stereoselective Analysis of Methadone and EDDP in Laboring Women and Neonates in Plasma and Dried Blood Spots and Association with Neonatal Abstinence Syndrome. American journal of perinatology, doi:10.1055/s-0040-1701505 (2020).

64 Deacon, R. M. Assessing nest building in mice. Nature protocols 1, 1117–1119, doi:10.1038/nprot.2006.170 (2006).

65 Berrendero, F. et al. Increase of morphine withdrawal in mice lacking A2a receptors and no changes in CB1/A2a double knockout mice. The European journal of neuroscience 17, 315–324 (2003).

66 Ward, P., Moss, H. G., Brown, T. R., Kalivas, P. & Jenkins, D. D. N-acetylcysteine mitigates acute opioid withdrawal behaviors and CNS oxidative stress in neonatal rats. Pediatric research, doi:10.1038/s41390-019-0728-6 (2020).

67 Hill, J. M., Lim, M. A. & Stone, M. M. in Neuropeptide Techniques 131–149 (Springer, 2008).

68 Jones, A. F. & Sheets, P. L. Sex-Specific Disruption of Distinct mPFC Inhibitory Neurons in Spared-Nerve Injury Model of Neuropathic Pain. Cell Rep 31, 107729, doi:10.1016/j.celrep.2020.107729 (2020).

69 Suter, B. A. et al. Ephus: multipurpose data acquisition software for neuroscience experiments. Front Neural Circuits 4, 100, doi:10.3389/fncir.2010.00100 (2010).

70 Cheriyan, J. & Sheets, P. L. Altered Excitability and Local Connectivity of mPFC-PAG Neurons in a Mouse Model of Neuropathic Pain. The Journal of neuroscience: the official journal of the Society for Neuroscience 38, 4829–4839, doi:10.1523/jneurosci.2731-17.2018 (2018).

