## Supplementary Tables and Figures for "Prenatal Methadone Exposure Disrupts Behavioral Development and Alters Motor Neuron Intrinsic Properties and Local Circuitry"

**Supplementary Table 1 Dam and Offspring Methadone and Metabolite Concentrations.**

| Plasma (ng/mL) | | | | | | |
|---|---|---|---|---|---|---|
| | Gestational Day 18 | | Postnatal Day 1 | | Postnatal Day 7 | |
| | Methadone | EDDP | Methadone | EDDP | Methadone | EDDP |
| Dam | 63.5 ± 11.3 | 74.6 ± 22.0 | 21.4 ± 6.3 | 38.8 ± 5.3 | 16.0 ± 2.2 | 40.4 ± 3.9 |
| Offspring | | | 0.5 ± 0.1 | 3.0 ± 0.5 | 0.5 ± 0.1 | 1.4 ± 0.4 |
| Brain (ng/g) | | | | | | |
| | Methadone | EDDP | Methadone | EDDP | Methadone | EDDP |
| Dam | 248.9 ± 54.1 | 16.8 ± 2.0 | 92.3 ± 17.0 | 16.9 ± 3.0 | 73.0 ± 10.7 | 15.9 ± 1.3 |
| Offspring | 2100.8 ± 237.6 | 27.6 ± 8.4 | 7.9 ± 0.6 | 1.8 ± 0.2 | 3.1 ± 0.3 | 0.01 ± 0.01 |
| Placental (ng/g) | | | | | | |
| Placental | 3862.1 ± 258.4 | 1124.0 ± 84.4 | | | | |

n=3 dams + their respective litters per timepoint; n=17-20 offspring samples at G18, n=15 offspring at P1, n=17-18 offspring at P7. All tissue and blood samples were collected 2.5 hours following the morning administration of methadone. All data are mean ± SEM.

EDDP: 2-ethylidene-1,5-dimethyl-3,3-diphenylpyrrolidine. The limit of quantification for methadone and EDDP detection was 0.1 ng/mL and 0.05 ng/mL in the plasma, respectively, and 0.08 ng/sample and 0.04 ng/sample of placenta and brain for both methadone and EDDP.

**Supplementary Table 2 Test Statistics for Volumetric MRI Analysis.**

| Volume of Interest | t ratio | df | p value |
|---|---|---|---|
| Striatum (R) | 1.58 | 20 | 0.13 |
| Striatum (L) | 0.572 | 20 | 0.57 |
| Cortex | 0.411 | 20 | 0.69 |
| Hippocampus (R) | 0.156 | 20 | 0.88 |
| Hippocampus (L) | 1.11 | 20 | 0.28 |
| Thalamus | 0.224 | 20 | 0.82 |
| Cerebellum | 0.267 | 20 | 0.79 |
| Basal Forebrain-Septum | 1.70 | 20 | 0.11 |
| Hypothalamus | 0.891 | 20 | 0.38 |
| Amygdala (R) | 0.319 | 20 | 0.75 |
| Amygdala (L) | 0.338 | 20 | 0.74 |
| Brain Stem | 2.04 | 20 | 0.055 |
| Superior Colliculus | 0.648 | 20 | 0.52 |
| Olfactory Bulb | 1.57 | 20 | 0.13 |
| Midbrain (R) | 0.746 | 20 | 0.46 |
| Midbrain (L) | 1.19 | 20 | 0.25 |
| Inferior Colliculus (L) | 0.301 | 20 | 0.77 |
| Inferior Colliculus (R) | 0.281 | 20 | 0.78 |

Unpaired t tests, n=11 (4M:7F) PME, 11 PSE (6M:5F) mice. See Supplementary Fig. 8 for visual representation of the data. *R,* Right; *L,* Left

**Supplementary Table 3 Intrinsic Properties of L5 M1 Neurons.**

| | PME-Female | PSE-Female | PME-Male | PSE-Female | F Statistics:<br>Interaction<br>Sex<br>Exposure |
|---|---|---|---|---|---|
| **Resting membrane potential (mV)** | -63.1 ± 1.12 | -64.0 ± 1.02 | -63.2 ± 1.62 | -63.7 ± 1.24 | F=0.0226, p=0.88<br>F=0.00564, p=0.94<br>F=0.277, p=0.60 |
| **Holding current (pA)** | -100 ± 16.6 | -81.2 ± 17.1 | -107 ± 27.3 | -86.9 ± 19.7 | F=0.000955, p=0.98<br>F=0.091, p=0.76<br>F=0.855, p=0.36 |
| **Input resistance (Mohm)** | 71.6 ± 3.15 | 86.6 ± 5.45 | 75.2 ± 5.74 | 79.9 ± 3.57 | F=1.35, p=0.25<br>F=0.117, p=0.73<br>**F =4.93, p=0.030*** |
| **Voltage Threshold (mV)** | -38.4 ± 0.75 | -37.2 ± 0.859 | -38.5 ± 0.900 | -39.1 ± 0.840 | F=1.06, p=0.31<br>F=1.31, p=0.26<br>F=0.118, p=0.73 |
| **Current Threshold (pA)** | 223 ± 14.5 | 196 ± 2.0 | 237 ± 21.5 | 230 ± 13.6 | F=0.354, p=0.55<br>F=2.04, p=0.16<br>F=1.02, p=0.32 |
| **AP half-width (milliseconds)** | 7.76 e$^{-4}$ ± 1.99 e$^{-5}$ | 8.22 e$^{-4}$ ± 1.90 e$^{-5}$ | 7.68 e$^{-4}$ ± 2.16 e$^{-5}$ | 7.50 e$^{-4}$ ± 1.67 e$^{-5}$ | F=2.28, p=0.14<br>F=3.56, p=0.063<br>F=0.436, p=0.51 |
| **Tau (milliseconds)** | 2.46 e$^{-3}$ ± 2.26 e$^{-4}$ | 2.46 e$^{-3}$ ± 1.86 e$^{-4}$ | 2.21 e$^{-3}$ ± 1.66 e$^{-4}$ | 2.21 e$^{-3}$ ± 2.20 e$^{-4}$ | F~0, p>0.99<br>F=0.111, p=0.74<br>F~0, p>0.99 |
| **FI Slope (Hz/pA)** | 0.082 ± 0.003 | 0.086 ± 0.004 | 0.081 ± 0.005 | 0.086 ± 0.002 | F=0.00942, p=0.92<br>F=0.0233, p=0.88<br>F=1.39, p=0.24 |
| **Voltage Sag (%)** | 49.1 ± 2.55 | 40.8 ± 3.09 | 46.3 ± 4.08 | 39.5 ± 3.37 | F=0.0507, p=0.82<br>F=0.379, p=0.54<br>**F=5.14, p=0.027*** |
| **Voltage Overshoot (%)** | 49.1 ± 2.27 | 41.1 ± 2.47 | 42.1 ± 3.31 | 39.5 ± 3.12 | F=0.881, p=0.35<br>F=2.24, p=0.14<br>F=3.40, p=0.070 |
| **Fast after-hyperpolarization (mV)** | -2.70 ± 0.562 | -3.63 ± 0.710 | -4.62 ± 1.24 | -3.20 ± 0.558 | F=2.15, p=0.15<br>F=0.863, p=0.36<br>F=0.0933, p=0.76 |
| **Medium after-hyperpolarization (mV)** | -12.0 ± 0.917 | -11.4 ± 0.939 | -7.48 ± 2.44 | -9.15 ± 1.00 | F=0.601, p=0.44<br>**F=5.50, p=0.022***<br>F=0.153, p=0.70 |
| **Afterdepolarization (mV)** | 24.4 ± 5.81 | 10.3 ± 6.28 | 12.1 ± 5.66 | 19.4 ± 7.81 | F=2.51, p=0.12<br>F0.0569, p=0.81<br>F=0.252, p=0.62 |
| **Height (mV)** | 83.7 ± 1.37 | 85.6 ± 0.983 | 86.9 ± 1.30) | 87.5 ± 1.65 | F=0.192, p=0.66<br>F=2.96, =0.090<br>F=0.711, p=0.40 |

Resting membrane potential evaluated with no applied current, all other properties evaluated with current applied to hold the membrane potential near minus 70 mV. Data presented as mean ± SEM. F statistics (df = 1,65) are presented in the final column with significant results bolded (*p<0.05). n=10 PME mice (6M:4F), 30 cells (13M:17F) and n=9 PSE mice (5M:4F), 23 cells (11M:12F).

**Supplementary Table 4 Antibody List.**

| Antibodies | Source | Identifier |
|---|---|---|
| Anti-Cux1 | Santa Cruz Biotechnology | Cat# SC-13024, RRID: AB_2261231 |
| Anti-NeuN | Millipore | Cat# MAB377, RRID: AB_2298772 |
| Goat anti-Mouse IgG (H+L), Alexa Fluor 488 conjugate | Thermo Fisher Scientific | Cat# A-11001, RRID: AB_2534069 |
| Goat anti-Rabbit IgG (H+L), Alexa Fluor 555 conjugate | Thermo Fisher Scientific | Cat# A-32732; RRID: AB_2633281 |
| Steptavidin, Alexa Fluor 488 | Thermo Fisher Scientific | Cat# S32354; RRID: RRID:AB_2315383 |

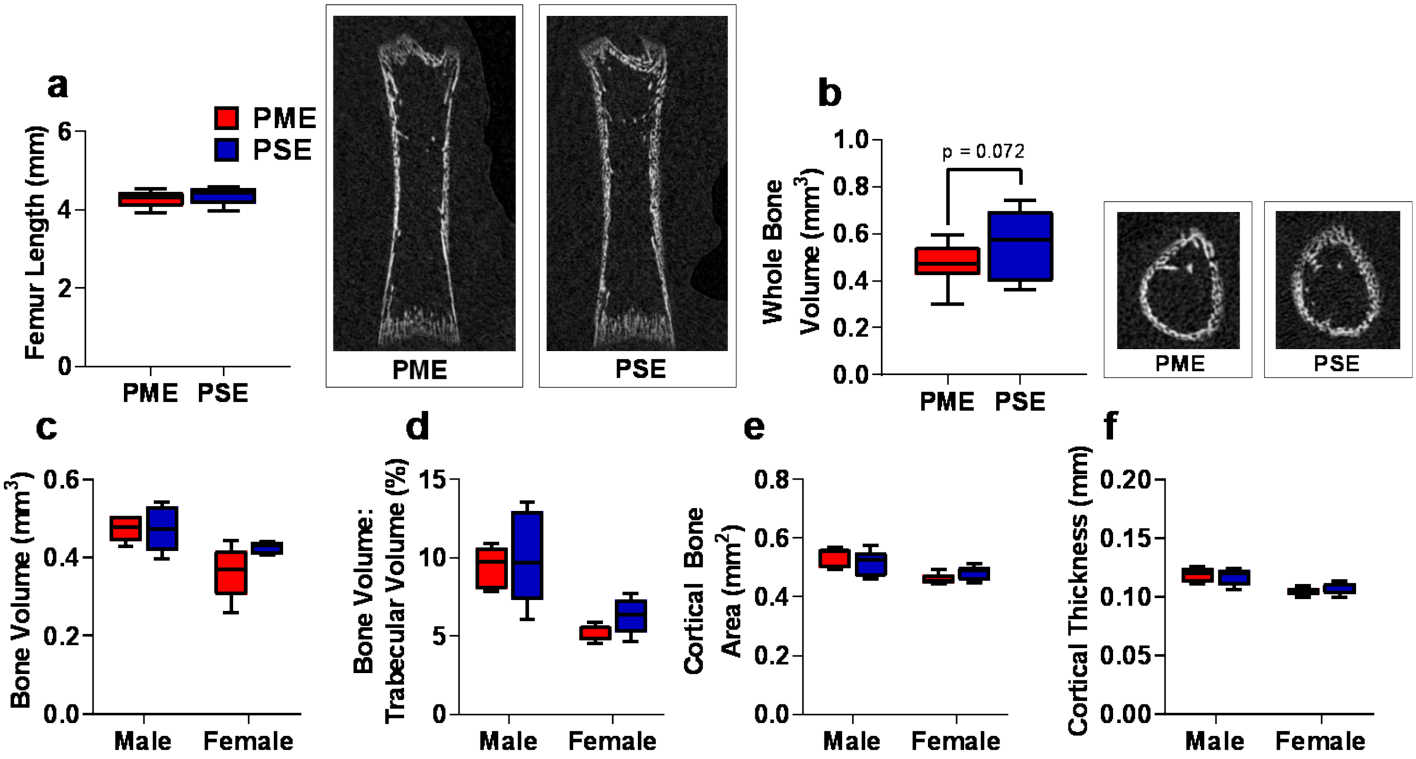


**Supplementary Figure 1 Early Life Bone Density May be Reduced in Prenatal Methadone-Exposed Offspring but Recovered by Adolescence. a** Femur length in P7 offspring did not differ between exposure groups. *Right* representative coronal CT image of P7 femur in PME and PSE offspring. **b** Femur whole bone volume was non-significantly reduced in P7 offspring with PME. *Right* representative CT image of P7 offspring demonstrating transaxial section of femur midshaft in PME and PSE offspring (unpaired t test: p=0.072, n=13 PME (6M:7F), 12 PSE (6M:6F) at P7). At P35, structural bone measures including **c** distal metaphysis bone volume, **d** trabecular bone volume, **e** cortical bone area, and **f** cortical thickness did not differ between exposure groups. (n=10 PME (5M:5F), 10 PSE (5M:5F) at P35). Box plots indicate 25th to 75th percentiles with whisker characterizing the minimum and maximum value.

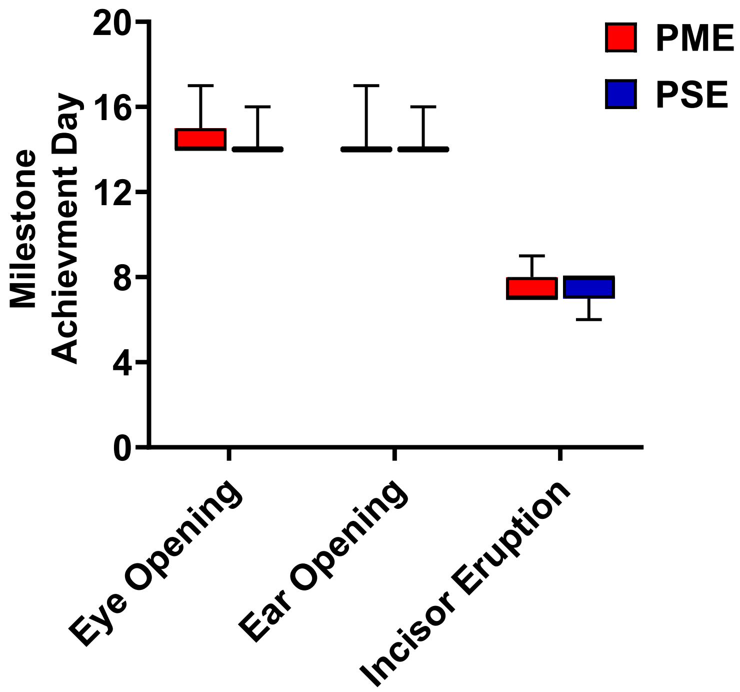


**Supplementary Figure 2 Postnatal Craniofacial Milestones in Offspring is Not Affected by PME.** Mice were examined daily to examine progress of craniofacial milestone measures. PME did not delay the day when both eyes opened, both ears moved to their final erect position and external auditory canals were patent, or when the bottom incisor teeth erupted (n=26 (7M:19F) PME, 25 PSE (15M:10F) mice). Box plots indicate 25th to 75th percentiles with whisker characterizing the minimum and maximum value.

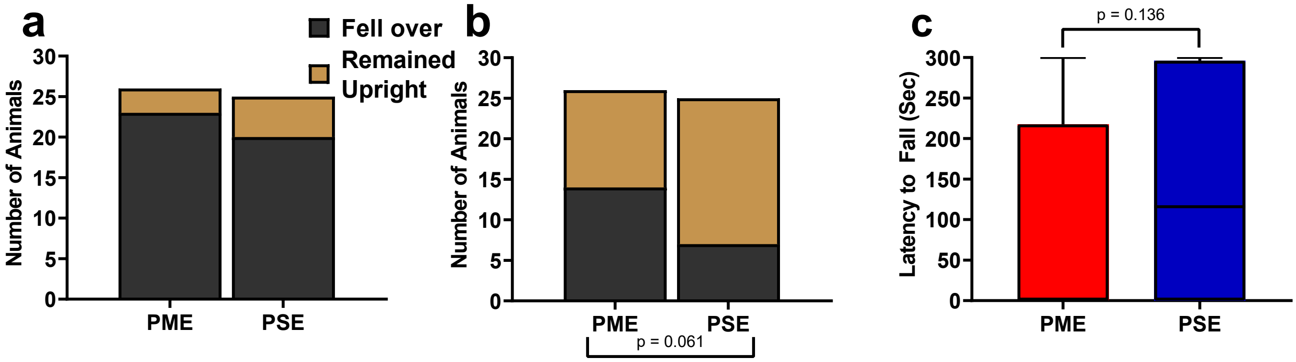


**Supplementary Figure 3 Prenatal Methadone Exposed Offspring Displayed a Greater Propensity to Fall Over on Postnatal Day 1. a** While the nearly all P1 offspring fell over by the end of the five-minute testing session (chi square test: p=0.41) **b** 14/26 PME compared to 7/25 PSE offspring had fallen over immediately upon placement into the arena (chi square test: p=0.061). **c** Similarly, PME offspring also display a nonsignificant reduction in the latency to fall over during the five-minute session (Mann-Whitney test: p=0.136 n=26 (7M:19F) PME, 25 PSE (15M:10F) mice. Bars indicate proportion of offspring that rolled over or remained upright. Box plots indicate 25th to 75th percentiles with whisker characterizing the minimum and maximum value.

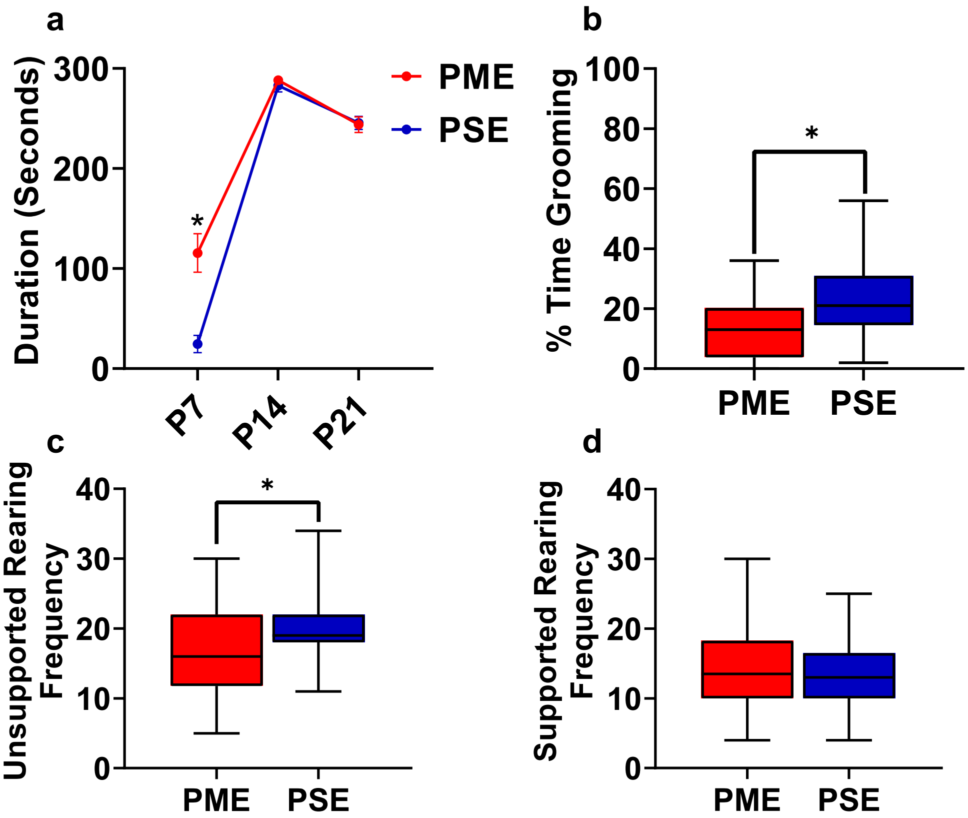


**Supplementary Figure 4 Open Field Behaviors Are Altered in Offspring With PME a** PME offspring spent significant more time in the open near the arena walls at P7 (rmANOVA: Exposure, p=0.0001; Interaction, p<0.0001; Sidak's post hoc test: p=0.0004), however no effect was observed on P14 or P21. As PSE animals were significantly less active at P7, it is likely that PSE animals did not leave the center of the arena where offspring were placed at the beginning of each trial. **b,c,d** Grooming and rearing were further assessed in P21 juvenile mice. **b** The total percentage of time grooming was significant reduced in PME mice (Mann Whitney test: p=0.0035). **c** PME offspring demonstrated significantly less unsupported rearing instances on P21 (unpaired t test: p=0.017), **d** but now differences in rearing supported by the arena walls was observed between exposure groups. (n=26 (7M:19F) PME, 25 PSE (15M:10F) mice). Data points indicate mean ±SEM and box plots indicate 25th to 75th percentiles with whisker characterizing the minimum and maximum value.

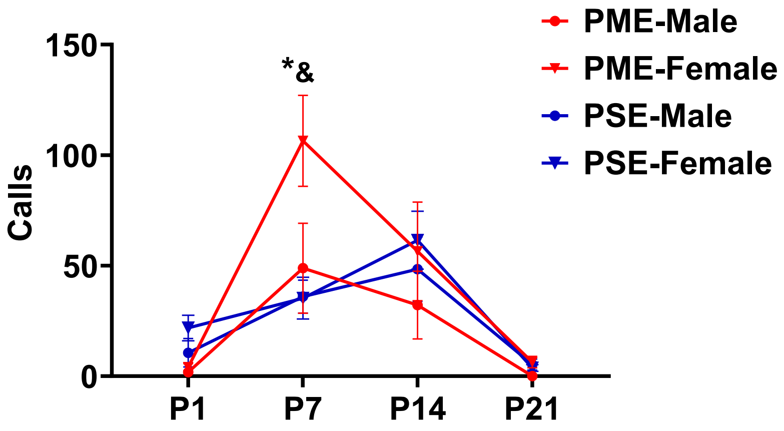


**Supplementary Figure 5 Separation Induced Ultrasonic Vocalizations Separated by Sex.** While there was a significant time by exposure interaction on the development of USVs (rmANOVA: p=0.032), sex effects were present on the production of USVs (rmANOVA: p=0.045). Although female PME did not vocalize more than male PME offspring on P7 (Sidak's post hoc test: p=0.60), female PME did emit more USVs than both male and female PSE offspring on P7 (Sidak's post hoc test: p=0.0045 & p=0.026, respectively, n=26 (7M:19F) PME, 25 PSE (15M:10F) mice). * p<0.05 for female PME vs female PSE; & p<0.05 for female PME vs male PSE offspring. Data points indicate mean ±SEM.

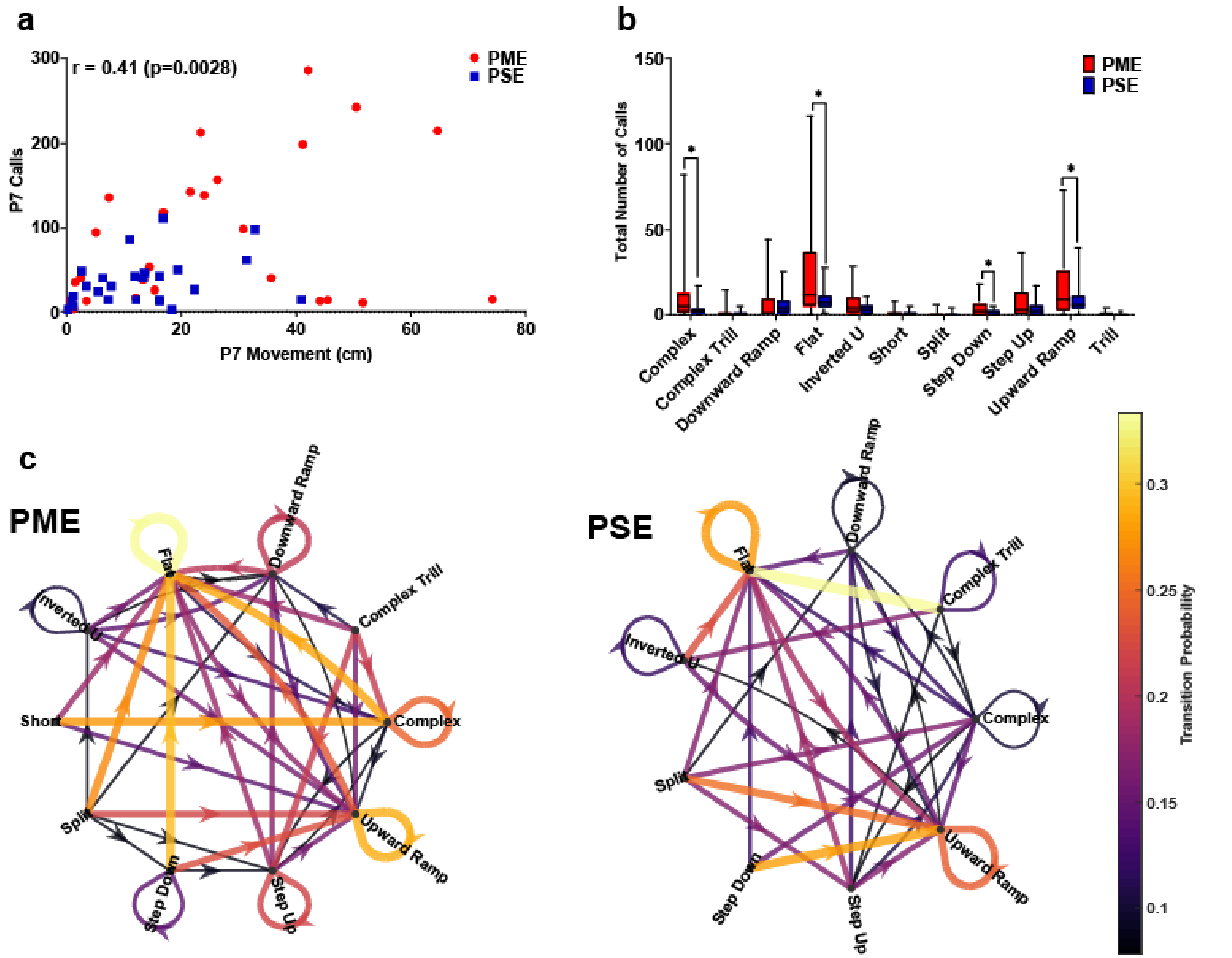


**Supplementary Figure 6 In Depth Analysis of Ultrasonic Vocalization Revealed Differential Patterns of Call Types and Syntax at P7. a** A moderate correlation exists between total locomotor activity and total USV calls at P7 suggesting a relationship is present between the hyperactivity and increased vocalizations among PME animals. **b** DeepSqueak automated classification of USV call types on P7 revealed an increase in production of all eleven call types in PME offspring but the level of significance was reached for complex, flat, step down, and upward ramp calls (unpaired t tests: p=0.014, p=0.0045, p=0.0086, & p=0.041, respectively). **c** Syntax analysis was also completed with DeepSqueak to determine the probably of call type transitions on P7 in both exposure groups. The probability of call transitions for PME and PSE offspring exhibit qualitative differences in syntax patterns suggesting alterations in maternal-pup communication may be present offspring with PME. Arrows represent the direction of call transition, and line thickness and color represent the transition probability (n=26 (7M:19F) PME, 25 PSE (15M:10F) mice). *p<0.05. Box plots indicate 25th to 75th percentiles with whisker characterizing the minimum and maximum value.

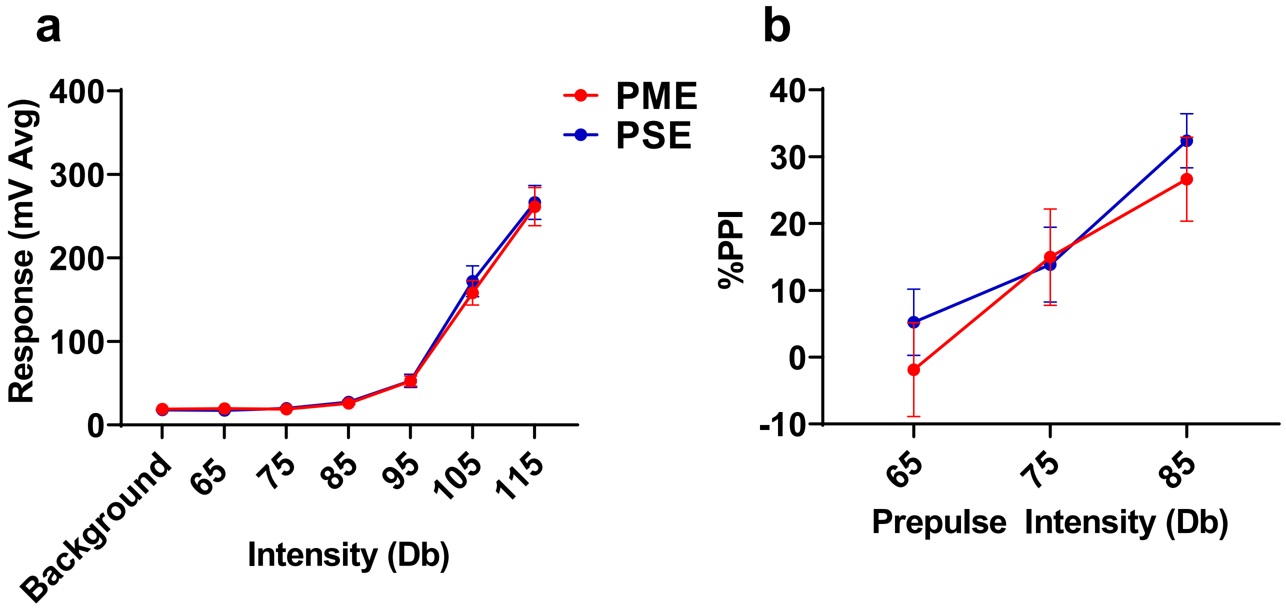


**Supplementary Figure 7 Acoustic Startle Response and Prepulse Inhibition was Not Affected by PME.** PME did not disrupt the **a** acoustic startle response to any startle intensity examined or **b** alter prepulse inhibition (PPI) at any prepulse intensity studied in P28-P29 offspring (n=19 (10M:9F) PME, 19 PSE (10M:9F) mice). Data points indicate mean ±SEM.

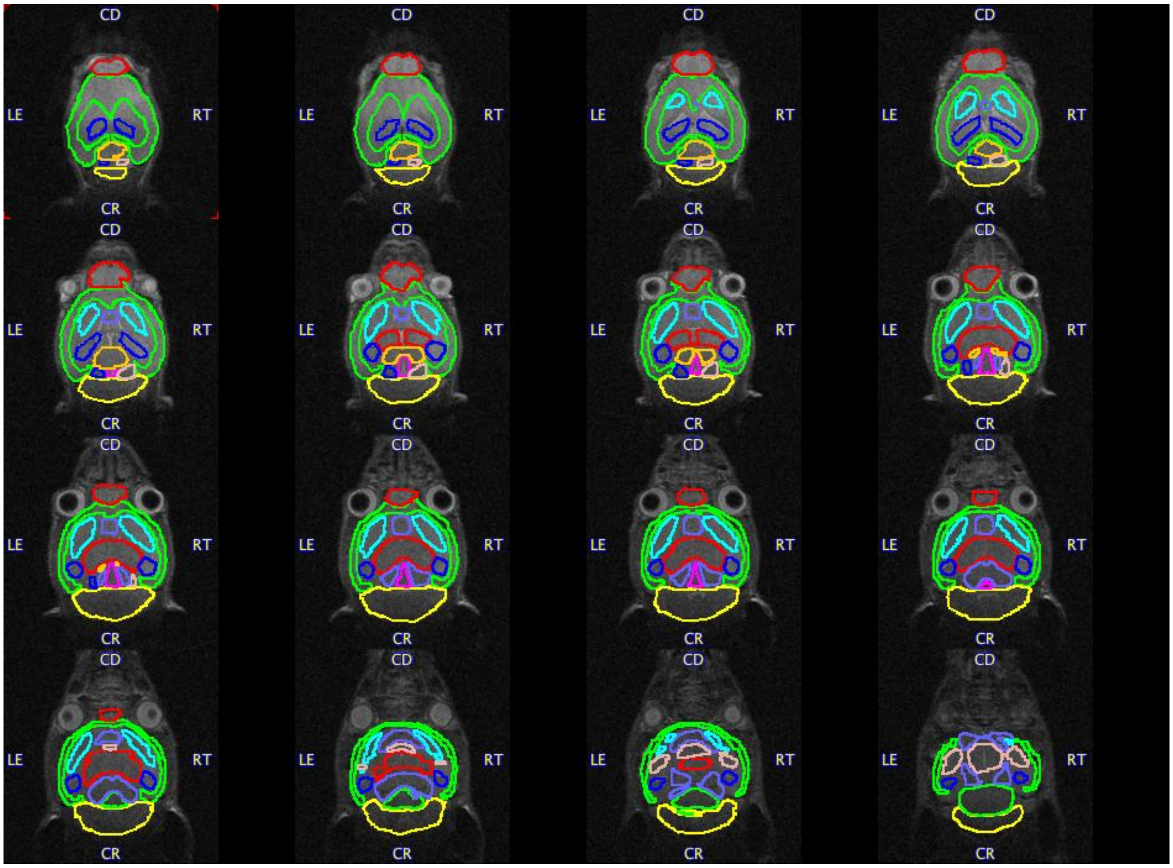


**Supplementary Figure 8 Brain Regions Examined for Volume Differences in Offspring.** Volumes of interest (VOIs) are overlaid on a representative mouse brain. A total of 18 VOIs segmented including the whole cortex, striatum (L & R), hippocampus (L & R), thalamus, cerebellum, basal-forebrain septum, hypothalamus, amygdala (L & R), brain stem, superior colliculus, olfactory bulbs, midbrain (L & R), and inferior colliculus (L & R). *R,* Right; *L,* Left.

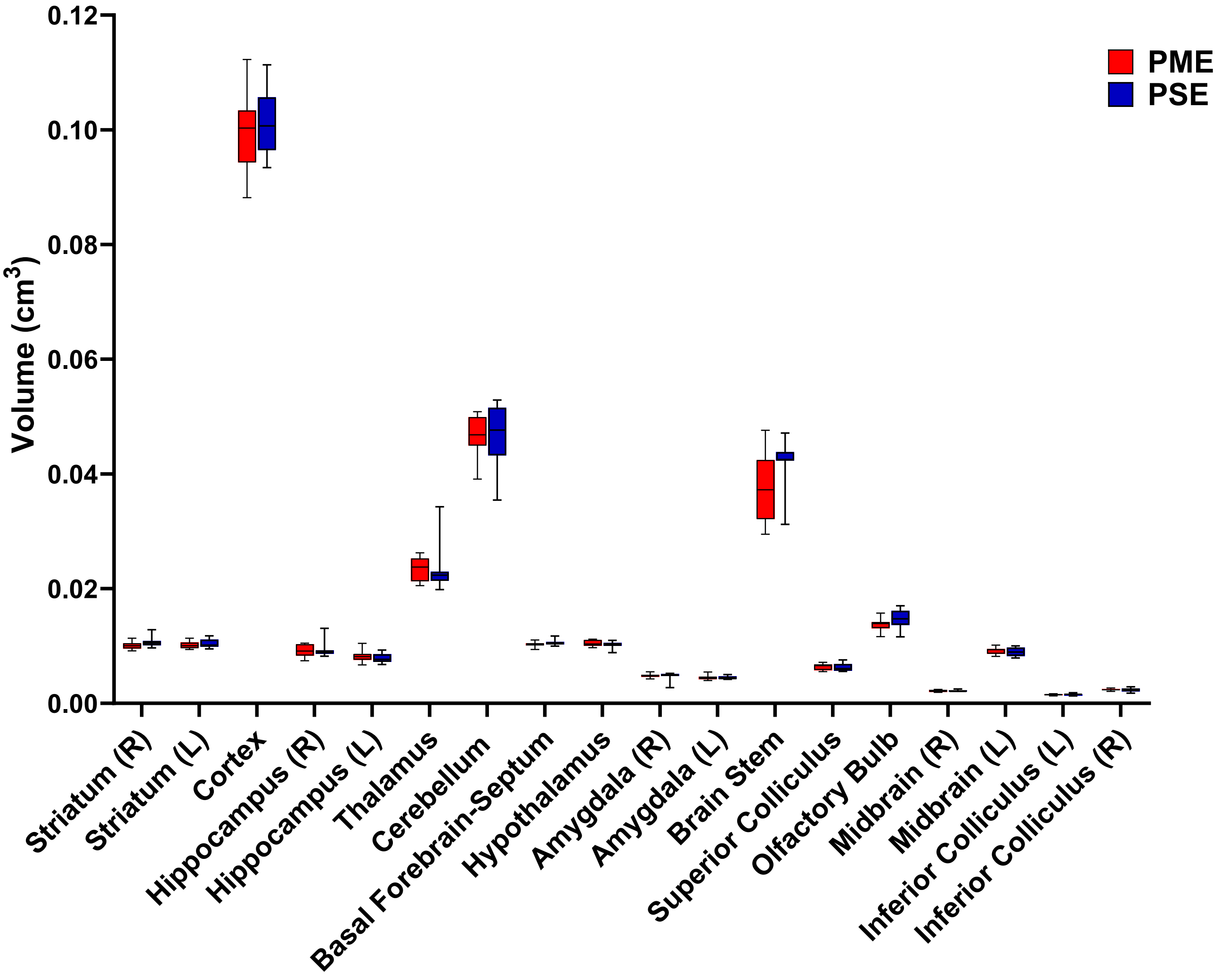


**Supplementary Figure 9 PME Did Not Reduce Volumes in Brain Regions of Interest.** Volumetric findings from ultra-high field MRI of P28-P30 offspring indicate that gross brain structure is primarily unaffected for both cortical and subcortical regions in PME offspring (unpaired t tests, n=11 (4M:7F) PME, 11 PSE (6M:5F) mice). Box plots indicate 25th to 75th percentiles with whisker characterizing the minimum and maximum value.

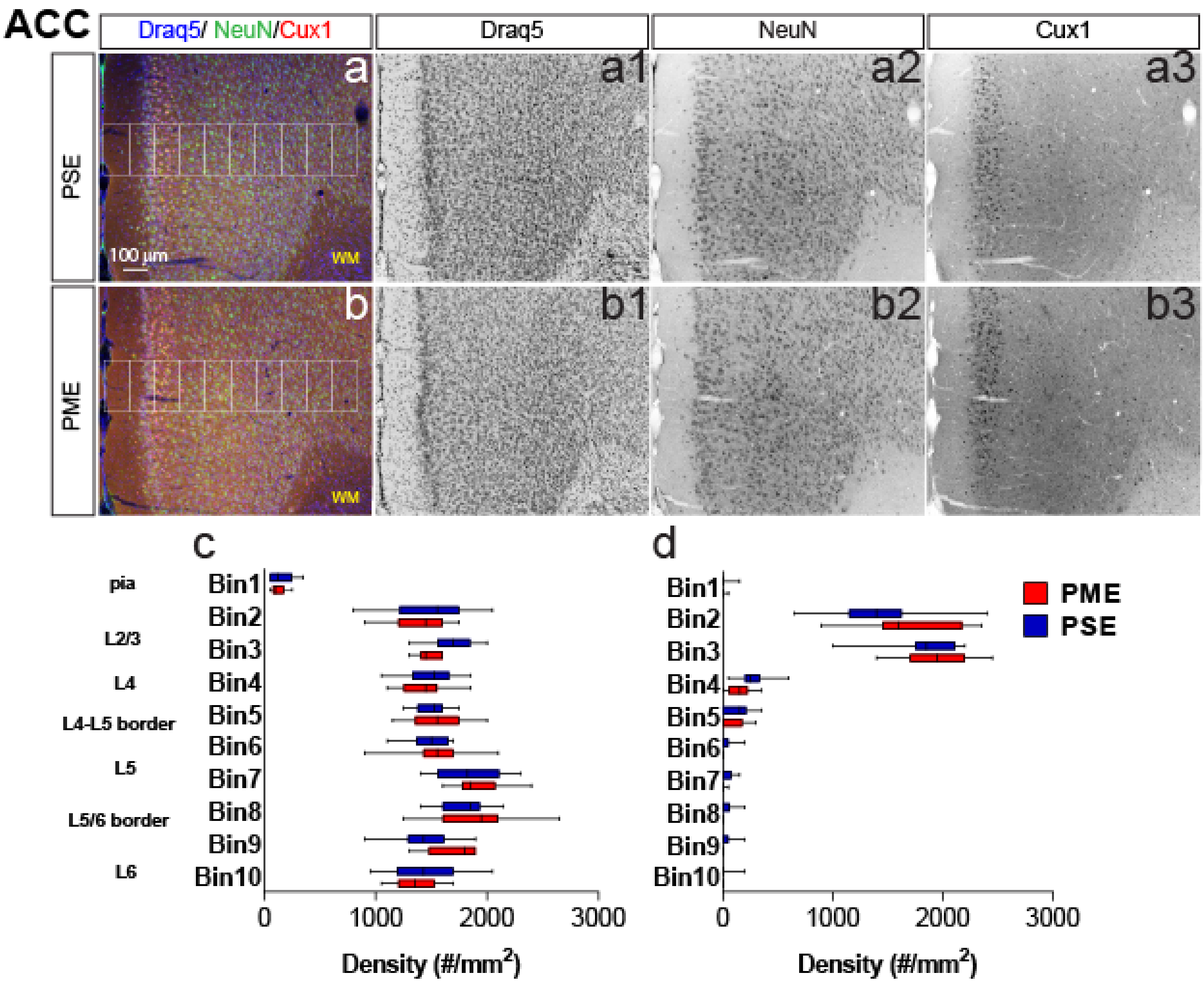


**Supplementary Figure 10 Development of the Anterior Cingulate Cortex (ACC) is Not Affected by PME.** Representative slices of the ACC in **a** PSE and **b** PME offspring demonstrating Draq5 (blue, a1,b1: marker of nuclei), NeuN (green, a2,b2,: marker of post-mitotic neurons), and Cux1 (red, a3,b3: marker of upper cortical layers, typically layer II-IV). White boxes represent bins 1-10 for quantification with Bin 1 nearest to the pia and Bin 10 nearest internal white matter. No effect of PME on **c** NeuN$^{+}$- or **d** Cux1$^{+}$-cell densities were observed in Bins 1-10 (n=7 (2M:5F) PME, 7 PSE (3M:4F). Box plots indicate 25th to 75th percentiles with whisker characterizing the minimum and maximum value

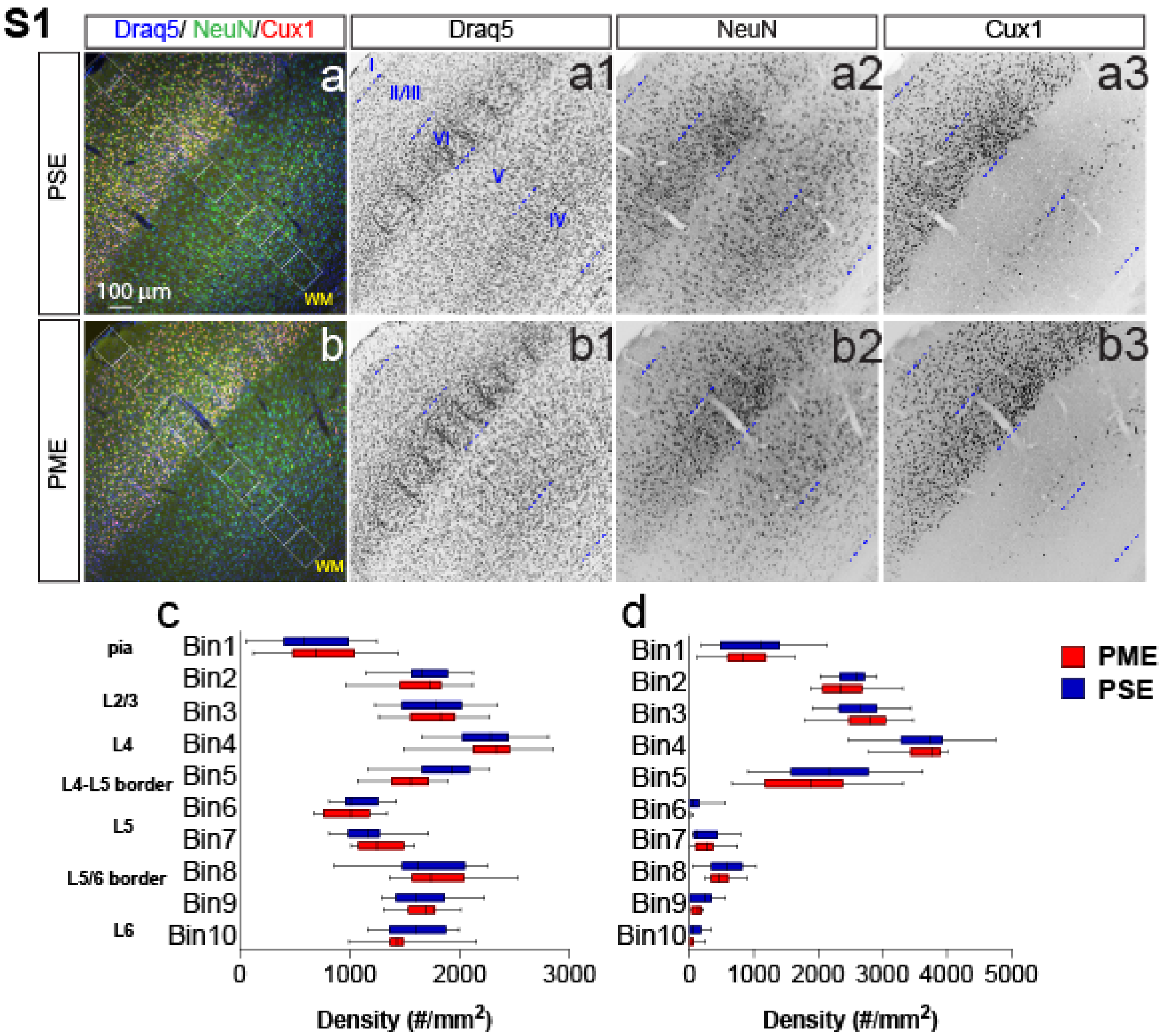


**Supplementary Figure 11 Development of the Primary Somatosensory Cortex (S1) is Not Affected by PME.** Representative slices of the S1 in **a** PSE and **b** PME offspring demonstrating Draq5 (blue, a1,b1: marker of nuclei), NeuN (green, a2,b2,: marker of post-mitotic neurons), and Cux1 (red, a3,b3: marker of upper cortical layers, typically layer II-IV). White boxes represent bins 1-10 for quantification with Bin 1 nearest to the pia and Bin 10 nearest internal white matter. No effect of PME on **c** NeuN$^+$- or **d** Cux1$^+$- cell densities were observed in Bins 1-10 (n=7 (2M:5F) PME, 7 PSE (3M:4F)). Box plots indicate 25th to 75th percentiles with whisker characterizing the minimum and maximum value.

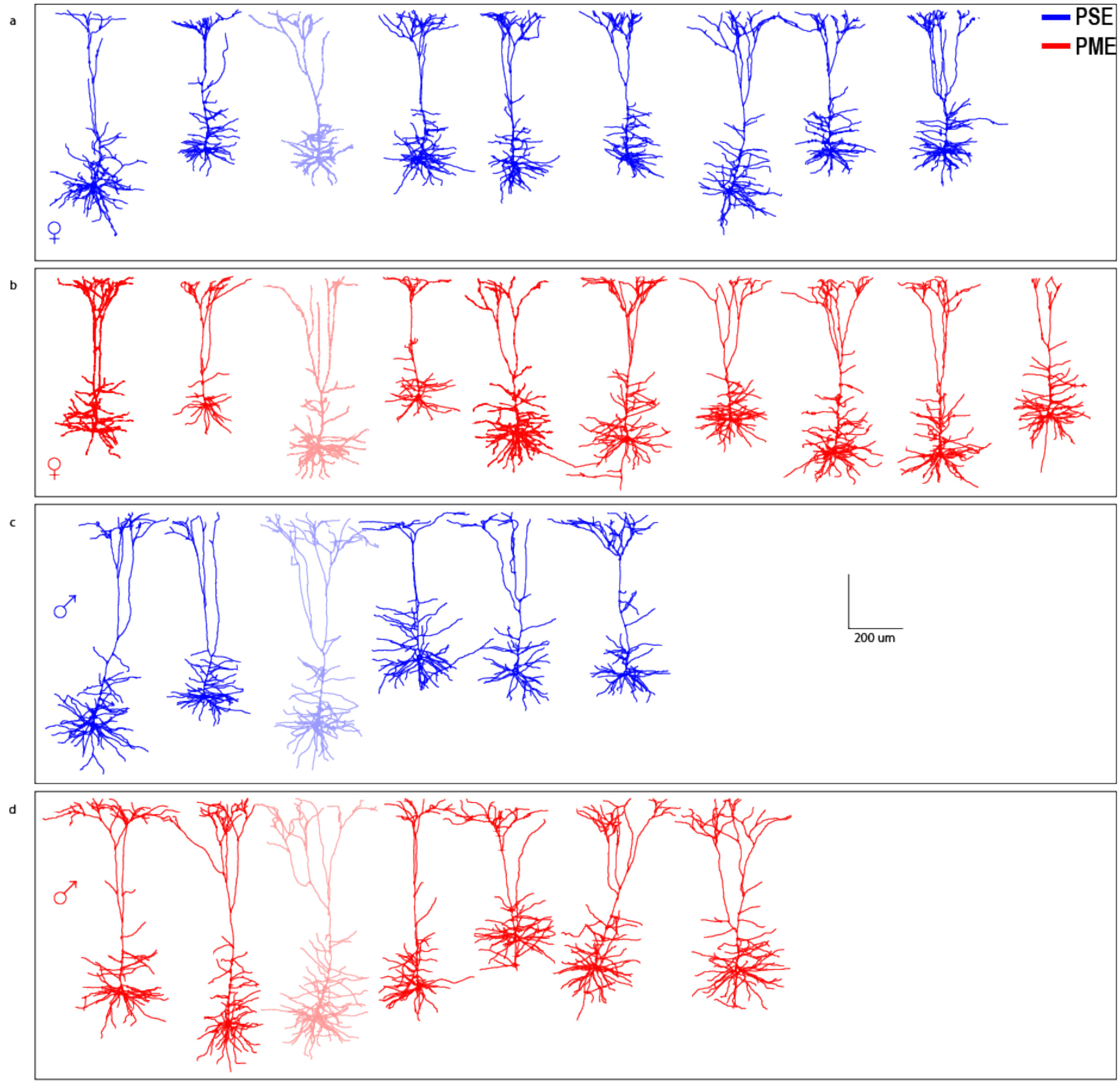


**Supplementary Figure 12 Reconstruction of M1 L5 Neurons in Offspring**
Morphological reconstructions confirm thick-tufted layer V M1 neurons recorded from a female PSE, b female PME, c male PSE, and d male PME mice. Partially opaque neurons are represented in Supplementary Fig. 14.

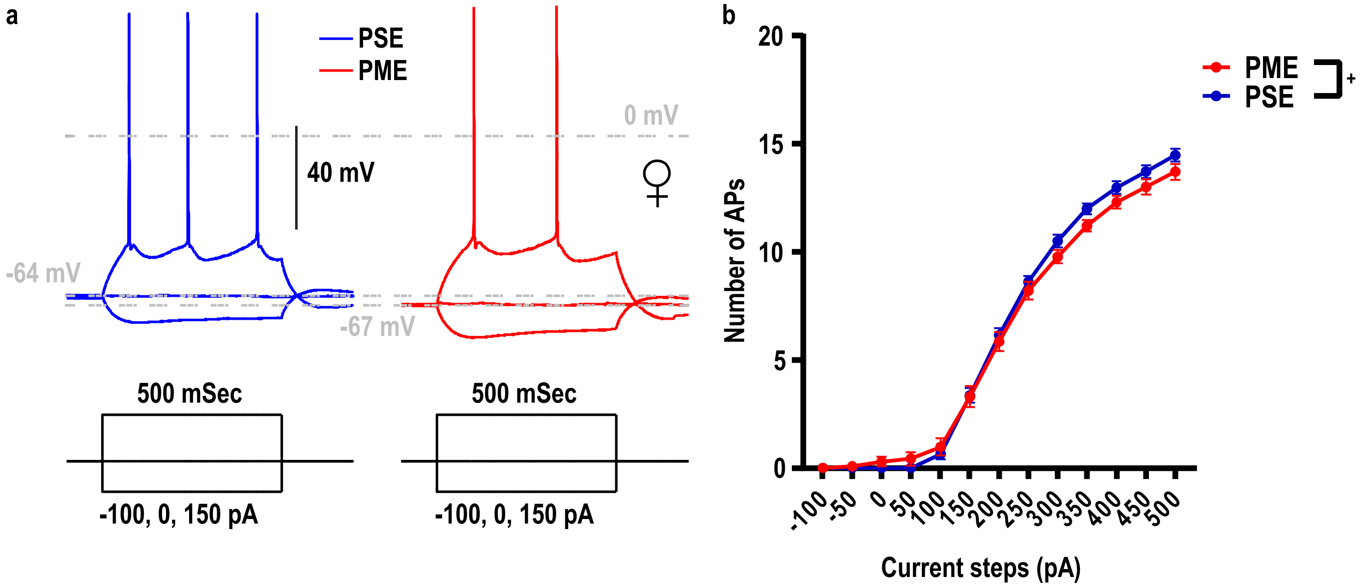


**Supplementary Figure 13 Current Threshold for Action Potential (AP) Firing in L5 M1 Neurons is Impacted by PME. a** Representative current-clamp traces of action potential firing from L5 motor cortex neurons from PSE (blue traces) and PME (red traces) female mice. **b** The number of action potentials in response to injected current in L5 M1 neurons revealed PME mice exhibit altered excitability (n=40 (15M:25F) PME, 28 PSE (15M:13F)). Data points indicate mean ±SEM. +: Current x Exposure Interaction, p=0.047.

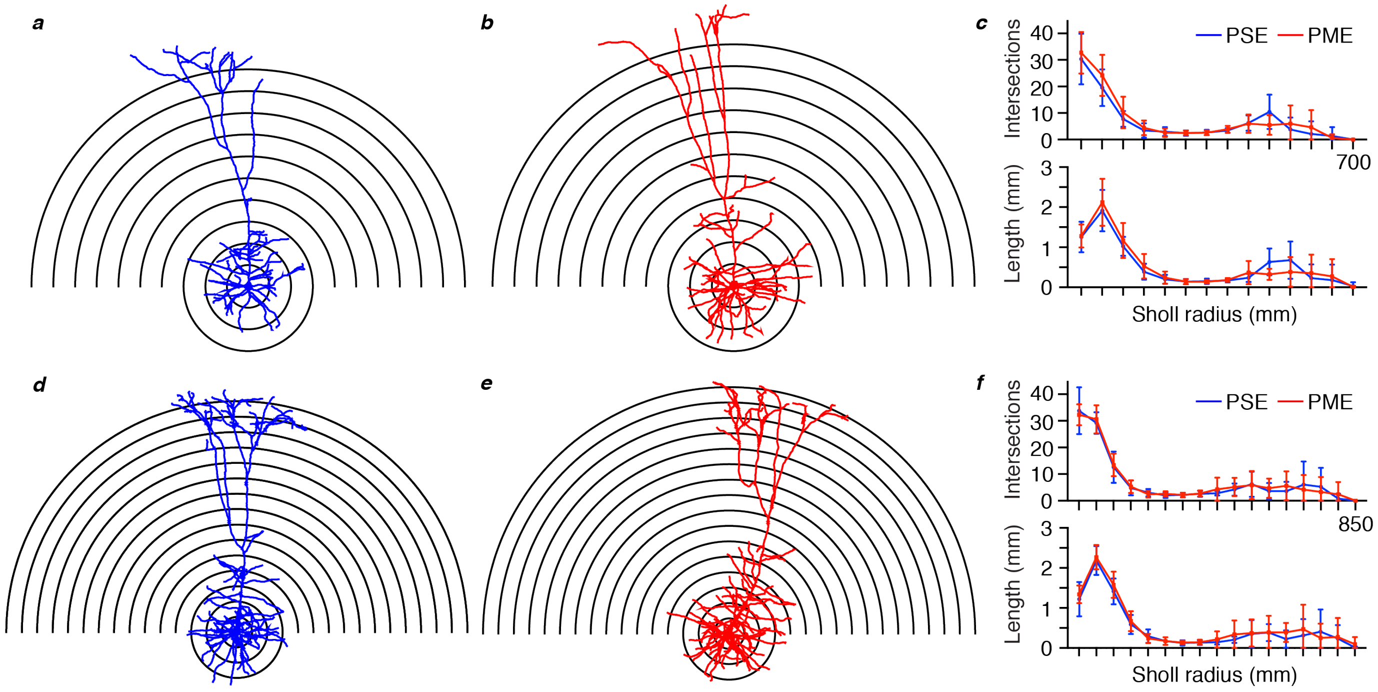


**Supplementary Figure 14 Morphological Analysis of L5 M1 Neurons Revealed No Effect of PME. a-b, d-e** Representative morphological reconstruction and Sholl radii of motor cortex neurons filled with biocytin during whole-cell patch-clamp recordings in acute brain slices from PSE and PME female **a-b** and male **d-e** mice (concentric circles increase by 50 µm from 50 µm). **c** Analysis of PSE (n=9) and PME (n=10) intersections (top) and length (mm, bottom) in female mice by Sholl radius (µm, 50:50:700) revealed no significant differences related to exposure. **f** Analysis of PSE (n=6) and PME (n=7) intersections (top) and length (mm, bottom) in male mice by Sholl radius (µm, 50:50:850) also revealed no significant differences related to exposure.
